# Social control, not service quality, explains fast growth in the cleaner wrasse *Labroides dimidiatus*

**DOI:** 10.64898/2026.05.16.725469

**Authors:** Letizia Pessina, Redouan Bshary

**Affiliations:** University of Neuchâtel, Faculté des sciences, Emile-Argand 11, 200 Neuchâtel, Switzerland

## Abstract

Interactions between cleaner fish *Labroides dimidiatus* and client fish, from which cleaners remove ectoparasites and mucus, represent a textbook example of mutualism involving sophisticated strategic decision-making. However, cleaners must also face intraspecific social challenges within a size-based hierarchy, where the largest females may eventually change sex and become males with higher reproductive rates. Following 540 individuals over 11 months, we found that, contrary to expectations, slow-growing females spent more time cleaning and cheated more frequently, without causing more negative client responses than fast-growing females did. Instead, variation in growth was best explained by social factors: fast-growing individuals experienced reduced social control, while slow growers spent more time in proximity to dominant individuals. As there was no evidence that spawning activity affected growth patterns, it appears that fast growth as a viable strategy for becoming a male largely depends on the lack of control by dominants.

## Introduction

Sequential hermaphroditism (sex change) may evolve when reproductive success differs between the sexes across body size or age, such that one sex achieves higher fitness when small or young and the other when large or old ^1–5^. Under these conditions, individuals can maximize lifetime reproductive success by functioning first as the sex with higher reproductive output at smaller sizes and later transitioning to the sex that achieves greater reproductive success at larger sizes ^5^. Because the terminal sex typically has the highest reproductive output ^5,6^, individuals of the initial sex may compete over access to the opportunity to change sex, raising the question of which features may enhance the likelihood of changing sex.

Protogyny, in which individuals begin life as females and later change into males ^1,5,7,8^, provides a useful framework for investigating how individuals navigate such competition. Protogyny is particularly common in haremic fishes, where a dominant male monopolizes reproduction within a group of females ^9^. In these systems, sex change allows a former female to take over a breeding territory, resulting in a substantial reproductive advantage ^5^. Because dominance hierarchies are typically size-based, the largest female, usually the highest-ranking individual, has the highest probability of changing sex and assuming the male role ^7–9,9–17^. This may create pressure on females to adopt strategies that increase their likelihood of reaching the terminal male stage.

Cleaner wrasse *Labroides dimidiatus* are a classic example of a monoandric protogynous hermaphrodite that lives in haremic social groups structured by size-based dominance hierarchies ^18–22^. All individuals are born female, and some large females may eventually change sex and become a harem male when the opportunity arises ^23^. This species is particularly interesting because males not only achieve higher reproductive output but also appear to have higher survival ^24^, making sex change a key life history objective for females in this system.

Recent work has further shown that these size-based hierarchies are unusually unstable compared with other well-studied systems, such as anemonefish (*Amphiprion spp.*) ^25–27^, coral gobies (*Paragobidon xanthosoma)* ^28^, or cooperative breeders like *Neolamprologus pulcher* ^29,30^, where the threat of eviction enforces strict social control and maintains highly stable rank structures. Apparently, the lack of cohesive group living, due to each female occupying her own core area, reduces the dominants’ control over subordinates and leads to common cases of rank reversal, in which smaller individuals outgrow initially larger harem members ^23^. These observations suggest that adopting a fast-growing life history could improve an individual’s future competitive position and its prospects of reaching the terminal male stage ^23^.

Currently, the mechanisms underlying variation in growth rate remain poorly understood. Here, we focus on a few key components linked to individual life-history decision-making, without investigating the underlying physiological mechanisms. Such key components are food intake, social control, mortality risk, and reproductive effort. Fast growers could be characterized by efficient/extended foraging, and/or increased mortality, and/or low investment in current reproduction, and/or reduced social control. In cleaner wrasses, foraging is a very interesting component of their lives. This is because nearly all food is obtained through cleaning interactions with client fishes ^31^, which visit cleaning stations to have ectoparasites removed ^31–35^. The interactions between the cleaners and clients represent a textbook example of mutualism involving sophisticated strategic decision-making. Although clients overall benefit from parasite removal ^36–39^, interactions are characterized by a conflict of interest: cleaners prefer to feed on client mucus rather than parasites ^40,41^, a behavior that constitutes cheating and negatively affects client health ^42,43^. Clients often respond to cleaner mucus feeding with a conspicuous body jolt, revealing the expressed conflict within this mutualism ^44^. In response, clients have evolved several behaviors that reduce a cheating cleaner’s gains and hence act as partner-control mechanisms, including punishment by resident species with access to the local cleaner only, partner switching by visiting clients with access to several cleaning stations within their home ranges, and image scoring ^44–51^. In turn, cleaners show high levels of strategic sophistication by adjusting the service quality to a variety of parameters, including client features such as size, parasite load, mucus quality, and maneuverability ^52,53^ as well as client strategic options ^40,51,54^. Individuals showing locally adaptive behavior harbor relatively larger forebrains than individuals failing to do so ^55^. Because cleaning interactions provide the primary source of food for cleaners ^56^, differences in time spent cleaning or in service quality could influence energy intake and, in turn, growth trajectories. Such factors might also be influenced by cleaner and client densities, which are known to differ between neighboring reefs within the same population and across years ^57,58^.

Variation in growth rates could also arise from individuals managing standard life history trade-offs ^59^ differently. Fast growth may cause increased mortality because of lowered somatic development ^60,61^, reduced immune function ^62,63^, as well as lowered resistance to physiological stressors ^64,65^. Non-mutually exclusive, investment in growth must be traded off against current reproduction ^66^.

A final factor that may influence growth rates concerns social interactions within the harem. In size-structured hierarchies, dominant individuals may suppress the growth of subordinates through aggression, thereby maintaining their competitive advantage ^25,27,28,30,67–73^. While dominant cleaner fish cannot threaten to evict smaller group members because each individual uses its unique core area ^23^, aggression and/or mere presence may still affect subordinate growth.

Here, we investigate factors that may contribute to fast growth as a correlate of achieving sex change in female cleaner wrasse. Specifically, we examine four main aspects: (i) time spent cleaning and service quality in their interactions with client fish, which may more globally be affected by cleaner and client densities, (ii) how service quality may relate to mortality risk, (iii) reproductive behavior, and (iv) social interactions with conspecifics. We expected that social competence expressed during cleaning interactions would be a main predictor of fast growth, meaning that we should identify an optimal level of cheating and providing tactile stimulation, which may depend on cleaner and client densities. More time spent cleaning should also translate into faster growth. Furthermore, we expected females showing less frequent evidence of reproductive activity to grow faster. Finally, individuals who are rarely targeted by larger, more dominant females should also grow fast, as low aggression and/or less time spent with dominants indicate a lack of social control. By linking growth trajectories with behavioral and ecological variables, we aim to identify how cleaning strategies, reproductive strategies, and dominance interactions interact to allow some females to increase their probability of reaching the terminal male stage.

## Methods

### Study site and population

Fieldwork was conducted at Lizard Island between July 2022 and June 2023 across eight reef sites (Supplementary Figure S1) differing in cleaner wrasse and client fish densities, as part of a long-term monitoring program. Full methodological details can be found in Pessina et al. (2026a,b) ^23,24^, as well as Supplementary Material S1. Individuals were captured via SCUBA using hand nets (10 x 15 cm) and barrier nets (4.7 x 1.8m or 1 x 1.2m), tagged in situ with visual implant elastomers (VIE), and released within two minutes to minimize disturbance. Adult fish received two-color tag combinations across multiple body positions (Supplementary Figure S2), while juveniles were marked with single high-contrast tags. Some individuals exhibited idiosyncratic features that enabled identification using natural markings (Supplementary Figure S3). In total, 540 individuals were tracked throughout the year (see Supplementary Table S1). Only the female phase of focal individuals was included in this study. Sex identification followed the methodology of Pessina et al. (2026a,b) ^23,24^, as detailed in Supplementary Material S1. Observations were conducted year-round (∼1,750 hours across ∼1,000 dives), allowing consistent individual recognition and monitoring. Tag retention and visibility remained high throughout the study, with no observed loss and confirmed durability through opportunistic recaptures. As sites were spatially isolated and cleaner wrasse exhibit strong site fidelity, inter-site movement was negligible, allowing independent site-based sampling.

### Transects to characterize cleaner and client densities

To quantify reef-scale ecological variables, including cleaner and client fish densities, we conducted standardized underwater visual census using belt transects. Fish counts included all reef fish, except for nocturnal and cryptic species. Surveys were conducted once during the Australian summer of 2023 and were stratified by the reef habitats where our focal cleaner resided: the reef crest and the reef base. The reef crest is defined as the outer edge of the reef flat ^74^, and the reef base as the lower reef slope where it meets the sandy substrate ^74^.

Within each habitat and site, five 30-meter transects, spaced 2 meters apart, were surveyed running parallel to the reef contour. Fish counts were completed running three consecutive swims per transect, each targeting a different size or taxonomic group: (i) large-bodied fishes swimming above the reef were counted within a 5 m wide belt; (ii) medium-sized fishes occupying the reef matrix were recorded within a 3 m belt; and (iii) damselfishes (Pomacentridae) were surveyed within a 2 m belt. The intermediate belt width was selected based on prior work demonstrating its suitability for accurately surveying wrasse species (Green 1994).

To minimize disturbance effects, each transect swim was preceded by a 2-minute pause. Transects were surveyed at a constant pace and completed in approximately 10 minutes. Only individuals measuring ≥4 cm total length were included in the analyses. Fish were identified to species level whenever possible, following WORMS taxonomic standards. Across all surveys, 343 species representing 51 families were recorded.

These counts were then used to calculate cleaner and client densities, which then allowed us to estimate cleaner-to-client ratios at our sites.

### Characterizing fast, slow, and average growers

Growth measurements were collected using an underwater stereo camera system following the approach of Pessina et al. (2026a,b) ^23,24^. A detailed methodological description is provided in Supplementary Materials S2. From these data, we reconstructed two types of growth curves: deme-specific growth trajectories and a population-level curve for Lizard Island.

The deme-specific curves were used to classify individuals into distinct growth strategies (fast, average, or slow) within each site. Classification was based on each fish’s final observed size relative to its deme-specific growth curve. Individuals exceeding half of the upper standard-deviation threshold were classified as fast growers, whereas those falling below half of the lower standard-deviation threshold were classified as slow growers; the remaining individuals were considered average growers. The use of half-standard-deviation thresholds was chosen to limit inflation of the average category while maintaining balanced group sizes.

These growth classifications were then used to examine differences in cleaning behavior, spawning frequency, and intraspecific interactions among growth strategies. In parallel, the population-level growth curve was used to assess how local cleaner and client densities influenced the distribution of growth strategies, allowing density effects to be evaluated independently of local growth dynamics.

### Collecting cleaning observations

Cleaning observations were collected for 34, 93, and 67 fast-, slow-, and average-growing adult focal females, respectively, totaling 914 videos (120 fast, 471 slow, 323 average). Behavioral observations consisted of 20-minute video recordings obtained using a GoPro Hero 9, with a scuba diver positioned approximately 2–3 m from the focal individual.

Each video was subsequently analyzed to characterize cleaning interactions. A cleaning interaction was defined as beginning with the first physical contact between the cleaner and a client and ending when the two individuals were separated for more than 5 seconds. For each interaction, we recorded: (i) client taxonomic identity (species when identifiable, otherwise genus); (ii) interaction duration; (iii) number of jolts; (iv) client responses to jolts, either through punishment of the cleaner or premature termination of the interaction; and (v) the occurrence of tactile stimulation.

Clients were assigned to one of two residency categories, namely resident or visitor. Because size information was unavailable for individual clients, classification was made at the family level (Supplementary Materials S3). Although most families can be classified as being mostly resident or visitor, we acknowledge the existence of exceptions (e. g., Pomacentridae is mostly resident, but the genus *Abudefduf* includes visitor-like species). Nevertheless, family-level classification remains a reasonable proxy for residency status. Moreover, given the greater complexity of the reef habitat compared to that studied by Bshary (2001) ^40^, applying a finer-scale methodology was not feasible.

From these observations, we constructed two datasets. The first dataset was used to address questions related to cleaning service quality. For each focal video, data were initially aggregated by client species cleaned during that recording. For each species, we calculated the total number of interactions, the total duration of cleaning, the total number of jolts, the percentage of jolts that elicited a negative response (punishment or termination of the interaction), and the percentage of interactions involving tactile stimulation. Jolt rate was calculated by dividing the total number of jolts by the total cleaning time. These metrics were then summarized at the video level by client type (resident or visitor), by summing the number of interactions and total cleaning time, and by averaging jolt rates, client responses to jolts, and tactile stimulation. Finally, data were aggregated to the cleaner ID level within each client type by averaging across all videos.

To examine client responsiveness to jolts, we constructed a second dataset following the same general workflow, with one key difference. To account for zero inflation in client responses, we did not perform an initial species-level aggregation. Instead, data were aggregated directly at the video and client-type levels.

### Collecting intraspecific observations

A total of 746 of those videos were also analyzed for intraspecific patterns (fast = 102, slow = 382, average = 262) for 177 focal cleaners (fast = 33, slow = 83, average = 61). Intraspecific analyses involved recording information of the interacting partner regarding their sex, relative size, and rank in relation to the focal individual, and identity when possible. The nature of cleaner-cleaner interaction varied from neutral to negative. Interactions involving either aggression towards or aggression received (through different degrees of chasing) from another cleaner were classified as negative interactions. When no negative interactions occurred in videos in which the focal cleaner encountered conspecifics, these instances were recorded as 0 and included in the analyses conducted in RStudio ^75^. Social rank for both focal and partner individuals was assigned using a monthly-updated social hierarchy dataset. To define social rank, individuals within each group were ordered strictly by body size (TL). *Labroides dimidiatus* is known to form a simple, stable, size-based dominance hierarchy governed by a “size principle,” in which aggression flows almost exclusively down the size gradient ^18,20–22,76^. We empirically validated this in our study population in Pessina et al. (2026a)^23^, where body size reliably indicated intra-group aggressive interactions, making the use of additional ranking algorithms based on behavioral matrices unnecessary. Using these data, we constructed a dataset to assess potential differences in the proportion of negative intraspecific interactions between growth strategies. For each Cleaner ID, we calculated the proportion of time during which the focal cleaner was either aggressing subordinates or being aggressed by dominants by dividing total interaction time invested in being attacked or attack by the sum of all videos’ durations. These proportions gave one value per fish. A similar methodology was used to build a dataset on the proportion of time each fish would spend in the presence of larger dominant individuals, without restricting to negative interactions.

From the same video analyses, a spawning dataset was also compiled. For each fish, three binary measures of reproductive behavior were recorded: (i) presence or absence of a distended belly, a proxy for egg development and the least precise but most abundant measure; (ii) female engagement in the Body-Sigmoid display, a behavior directly implying the presence of eggs ^21^; and (iii) actual spawning events, which necessarily imply the occurrence of the previous two behaviors. Because spawning activity in the cleaner wrasse is known to be influenced by lunar phase and tides ^18,21,77,78^, these variables were also included in the dataset. For each video, the lunar phase was classified as waxing, waning, full, or new. In addition, for tidal conditions, the temporal distance (in hours) between the video recording time and the nearest high tide was calculated. Only high tides occurring between sunrise and sunset were considered, as fish are largely inactive outside daylight hours. Videos that did not properly record the time of day in which they were taken were not considered, leading to a total of 373 videos (fast = 87, slow = 286, average = 286) for 177 cleaner fish focal individuals (fast = 32, slow = 83, average = 62).

### Statistical methods

Data analyses were conducted in R version 4.3.1 ^79^ using RStudio ^75^. Depending on the question, we used Generalized Linear Mixed-Effects Models (GLMM), Linear Mixed-Effects Models (LMM), Linear Models (LM), General Linear Models (GLM), and Pearson’s product–moment correlation tests. Models were fitted using lme4 ^80^, glmmTMB ^81^, and stats ^79^ packages. Model assumptions were verified through residual analyses and visual inspections of model fit. When appropriate, post hoc comparisons were performed with the emmeans package ^82^. Models were simplified based on non-significance and AIC values.

#### 1. Analyses related to cleaning behaviour

To test for differences in cleaning patterns between growth strategies, we fitted three LMMs and two GLMMs. All models include growth strategy, client type, site, and their interaction as fixed factors, and account for non-independence of observations by including a random intercept for cleaner ID.

First, we examined whether fast, slow, and average growers differed in the average time spent cleaning during a 20-minute video (Model 1). The response variable was log-transformed average cleaning duration (ln(duration + 3.05)), and the model was fitted with an LMM. The model accounted for heterogeneity in variability across growth strategies by modeling the dispersion parameter as a function of strategy, client type, and site.

Second, we tested for differences in the average number of cleaning interactions per video between growth strategies (Model 2). The response variable was the log-transformed average number of interactions (ln(number + 2.9)), and the model was fitted using an LMM.

The third and fourth models focused on investigating the quality of the cleaning service. The LMM model investigating potential differences in average jolt rate (Model 3) used the log-transformed (ln(response + 1.61)) as the response variable. The model investigating differences in average percentage of tactile stimulation was a GLMM that accounted for heterogeneity in variability across growth strategies by modeling the dispersion parameter of the beta distribution as a function of strategy, client type, and site (Model 4). Here, the average percentage of interactions with tactile stimulation was used as the response variable.

Using the second cleaning dataset, we then examined potential differences in client responsiveness to jolts between growth strategies (Model 5). This analysis was performed using a GLMM with a Tweedie distribution, with the response variable being the average proportion of jolts that were either punished or led to termination of the interaction.

In addition to these analyses, we investigated a potential link between service quality, growth strategy, and survival. To do so, we first performed a principal component analysis (PCA) using *dudi.PCA* function ^83^ on our centered and standardized cleaning variables. For this analysis, data were aggregated per cleaner ID by averaging per 20 minute video the total number of interactions (Tot int.), the total duration (Duration), the percentage of interaction with jolt (jolt), the percentage of interaction with tactile stimulation (TS), and the percentage of interactions with jolts that were negatively responded to (Resp.). Here, behavioral variables were aggregated across client types (i.e., resident and visitor clients were not analyzed separately) to avoid instability and dispersion issues in the model. We retained the first two principal components (PC1 and PC2), which together explained 63% of total behavioral variance (PC1 = 41%, PC2 = 22%). Individual PC scores were used in subsequent analyses.

To test whether survival probability was associated with cleaning behavior and growth strategy, we fitted a GLMER with a binomial distribution and a logit link function (Model 6). Here, the binary response variable was survival, PC1, PC2, growth strategy, and its interactions with the principal components were used as fixed factors, and site was included as a random intercept to account for spatial clustering and shared environmental conditions.

#### 2. Analyses related to cleaner and client densities

To examine associations between behavioral variables and environmental context, Pearson’s product–moment correlation tests were conducted between the proportion of time spent interacting with larger individuals, the proportion of time spent being aggressed by larger individuals, and cleaner reef fish density. Correlations were computed using complete-case observations (Models 7.1-7.2)

A GLM model with a binomial distribution was used to investigate whether the proportion of growth strategies is determined by the cleaner-to-client ratio (Model 8.1). The model included growth strategy, cleaner-to-client ratio, and their interaction as predictors. The model was fitted using the total number of fast and slow growers at each reef as weights. Pairwise correlations between cleaner density, client density, and cleaner-to-client ratio were computed using Pearson’s product-moment correlation tests as well (Model 8.2-8.4).

To examine associations between behavioral variables and environmental context, Pearson’s product–moment correlation tests were conducted between the proportion of time spent interacting with larger individuals, the proportion of time spent being aggressed by larger individuals, and cleaner reef fish density. Correlations were computed using complete-case observations (Models 7.1-7.2)

#### 3. Analyses related to spawning behavior

We computed three separate GLMMs models with a binomial response using the reproductive measures Belly (Model 9), Body-Sigmoid display (Model 10), and actual Spawning (Model 11) to investigate our question. Fixed effects included growth strategy and the distance to the nearest high tide, modeled with both linear and quadratic terms to capture non-linear patterns in spawning activity. The quadratic term was included to represent biologically realistic peaks in spawning probability at intermediate tidal distances, allowing estimation of the optimal distance for peak spawning using the formula 𝑥_peak_ = −𝛽_linear_/(2𝛽_quadratic_), where 𝛽_linear_and 𝛽_quadratic_are the coefficients of the linear and quadratic tidal terms, respectively. Random intercepts were included for individual fish to account for repeated observations, and for moon phase to account for phase-specific variation in spawning activity. Model fit and assumptions were assessed using residual diagnostics, and marginal and conditional R² values were calculated to quantify the proportion of variance explained by fixed effects alone and by both fixed and random effects.

#### 4. Analyses related to intraspecific interactions

We analyzed variation in intraspecific aggression using an LMM (Model 12). Specifically, we modelled the proportion of time cleaners spent engaged in negative interactions (i.e., attacking subordinates or being attacked by dominants) as a function of growth strategy, aggression type (attack vs. attacked), and their interaction. The response variable was log-transformed (ln(prop)) to improve normality and model fit, and models were fitted with a Gaussian error structure. To account for repeated observations, a cleaner identity nested within the site was included as a random intercept. Observations were weighted by the number of videos contributing to each estimate. Sample sizes were: fast (n = 66), average (n = 122), and slow (n = 166).

We then examined differences in the overall proportion of time spent interacting with larger individuals (irrespective of interaction type) using an additional LMM (Model 13). The log-transformed proportion of time (ln(prop)) was used as the response variable, with growth strategy as a fixed effect. Site was included as a random intercept to account for spatial clustering, and observations were weighted by the number of videos. Sample sizes were: fast (n = 33), average (n = 61), and slow (n = 83).

Cleaner reef fish density was not included in the main models because growth strategies were defined at the deme level using site-specific growth curves. Including density would have required using the categories at a population level, thereby losing the within-site resolution necessary to assess interaction patterns.

## Results

### 1. Cleaning patterns differ between growth strategies

Growth strategy and client type both strongly shaped cleaning behavior (Figure 1A). Cleaners differed in the average time spent cleaning depending on their growth strategy (Model 1; Analysis of Deviance: Chisq = 15.16, df = 2, p < 0.001) and the type of client serviced (Model 1; Analysis of Deviance: Chisq = 214.55, df = 1, p < 0.0001), with no significant interaction between the two predictors (Model 1; Analysis of Deviance: Chisq = 1.12, df = 2, p = 0.57). Site had an overall effect on cleaning duration (Model 1; Analysis of Deviance: Chisq = 21.41, df = 7, p = 0.003), but it did not interact with the effect of growth strategy (Model 1; Analysis of Deviance: Chisq = 16.48, df = 14, p = 0.29). Site did, however, interact with the effect of client type (Model 1; Analysis of Deviance: Chisq = 27.8, df=7, p < 0.0001; Supplementary Figure S6). Simplification of the model by removing non-significant interactions did not alter the significance of the main effects, and post hoc comparisons revealed that slow growers spent more time cleaning than fast growers (p = 0.0013; 57.5% and 52.6% more with resident and visitor respectively) but no difference was found for neither fast (p = 0.13) nor slow growers (p-value = 0.18) with average growers. Additionally, visitor clients received longer cleaning interaction than resident clients in all but two sites (p: 6 sites < 0.001; p: 2 sites > 0.05; Supplementary Figure S6).

**Figure 1:**
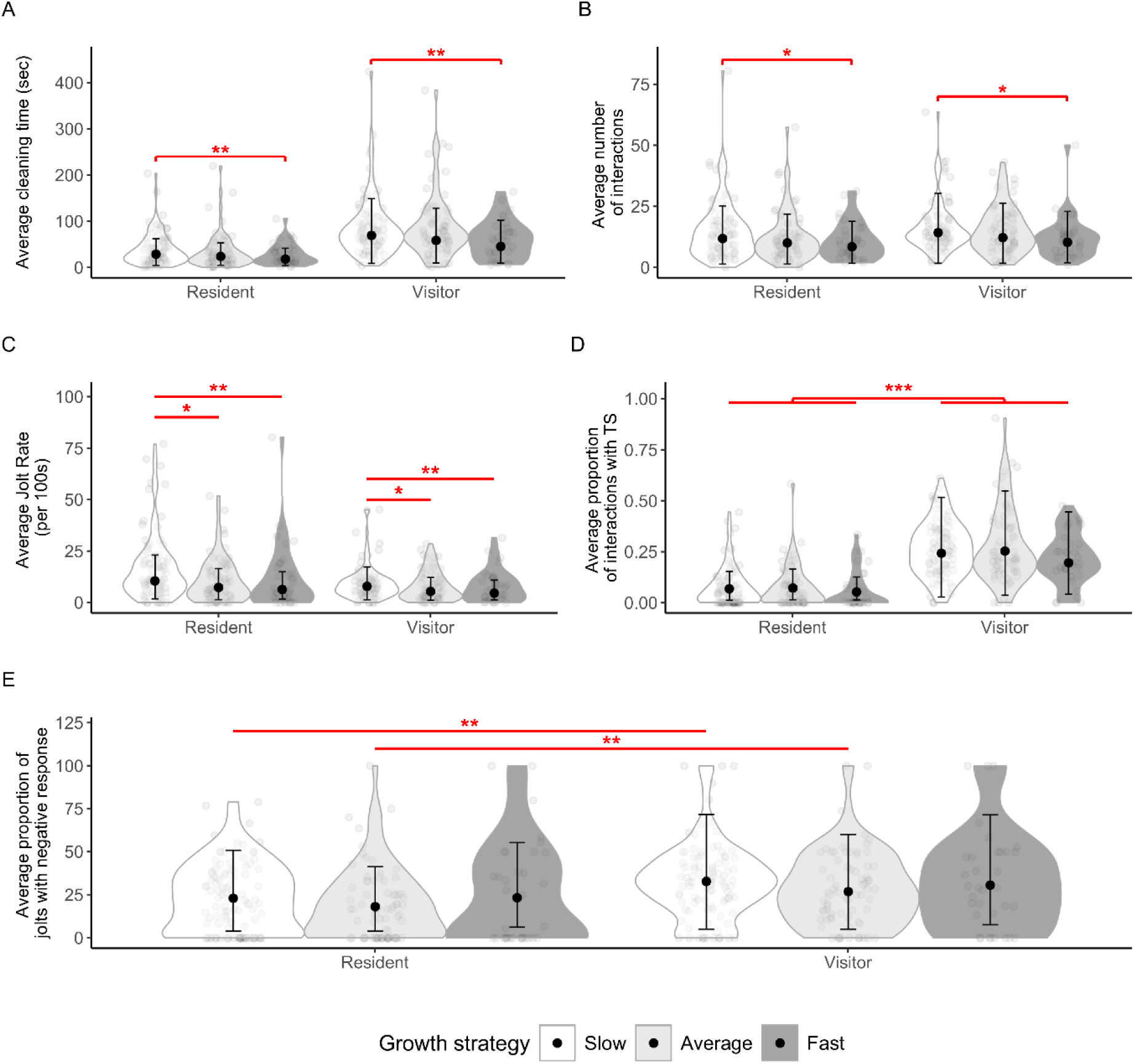
Cleaning patterns of fast, slow, and average growers. For each level of client type (resident and visitor) and growth strategy (fast, slow, and average), these graphs show: *A.* Distribution of average cleaning duration (Model 1; Gaussian LMM); *B*. Distribution of average number of cleaning interactions (Model 2; Gaussian LMM); *C.* Distribution of average jolt rate (Model 3; Gaussian LMM); *D.* Distribution of the average proportion of interactions with tactile stimulation (Model 4: Beta GLMM); and, *E.* Distribution of the average proportion of jolts with negative response (Model 5; Tweedie GLMM). All graphs show distribution as violin plots with jittered points representing raw individual-level data. The overlaid points and error bars indicate model-estimated means ± standard errors (SE). The line connecting the violins with an asterisk indicates a significant difference between the two groups involved (* *p* < 0.05; ** p < 0.01; *** p < 0.0001). Cleaning data includes 914 videos (fast = 120, slow = 471, average = 323) for 194 cleaner ID (fast = 34, slow = 93, average = 67).

Patterns in the number of interactions mirrored those observed for cleaning duration (Figure 1B). Both growth strategy (Model 2; Analysis of deviance: Chisq = 8.59, df = 2, p = 0.01) and client type (Model 2; Analysis of deviance: Chisq = 18.87, df = 1, p <0.0001) significantly affected the number of cleaning interactions, with no evidence for an interaction between predictors (Model 2; Analysis of deviance: Chisq = 2.29, df = 2, p = 0.32). Site was a significant predictor (Model 2; Analysis of deviance: Chisq = 14.87, df = 7, p = 0.04), but it did not significantly interact with the effect of growth strategy (Model 2; Analysis of deviance: Chisq = 13.06, df = 14, p = 0.52). However, Site did interact with client type (Model 2; Analysis of deviance: Chisq = 67.05, df = 7, p < 0.0001; Supplementary Figure S8). After model simplification, significance remained, and post hoc analyses showed that slow growers interacted more than fast growers (p = 0.02; 39.9% and 38% more with resident and visitor, respectively), but no difference was found for either fast (p = 0.35) or slow growers (p-value = 0.19) with average growers. Only at two of the study sites were visitor clients involved in significantly more interactions than residents (p: 6 sites > 0.05; 2 sites < 0.001, Supplementary Figure S8).

Cheating rates differed significantly between growth strategies (Model 3; Analysis of deviance: Chisq = 13.17, df = 2, p = 0.001, Figure 1C) and client types (Model 3; Analysis of deviance: Chisq = 9.64, df = 1, p = 0.002), while the interaction between growth strategy and client type was not statistically significant (Model 3; Analysis of deviance: Chisq = 2.38, df = 2, p = 0.30). Site had no significant effect on jolt rate (Model 3; Analysis of deviance: Chisq = 5.14, df = 7, p = 0.64) and it not interact with growth strategy (Model 3; Analysis of deviance: Chisq = 18.7, df = 14, p = 0.18), nor with client type (Model 3; Analysis of deviance: Chisq = 10.79, df = 7, p = 0.15). After model simplification, significances remained and post hoc analyses revealed that slow growers cheated more than both fast (p = 0.01; 65.6% and 45.3% more with resident and visitor respectively) and average growers (p = 0.02; 42.5% and 70.7% more with resident and visitor respectively), while no difference was found between fast and average growers (p = 0.67). Resident clients were cheated on more than visitor clients (p = 0.002).

Tactile stimulation showed a different pattern (Figure 1D). The proportion of interactions involving tactile stimulation depended strongly on client type (Model 4; Analysis of Deviance: Chisq = 223.02, df = 1, p < 0.0001), and marginally on site (Model 4; Analysis of Deviance: Chisq = 14.14, df = 7, p = 0.05) but was unaffected by growth strategy (Model 4; Analysis of Deviance: Chisq = 3.98, df = 2, p = 0.14). The interaction between growth strategy and client type was not significant (Model 4; Analysis of Deviance: Chisq =1.27, df = 2, p = 0.53). Similarly, site did not interact either with growth strategy (Model 4; Analysis of Deviance: Chisq = 16.48, df = 14, p = 0.28) or client type (Model 4; Analysis of Deviance: Chisq = 6.02, df = 7, p = 0.53). After model simplification, significances remained, and post hoc comparisons confirmed that cleaners provided more tactile stimulation to visitor clients than to resident clients (p < 0.0001).

Client responses to cheating were likewise primarily driven by client type (Figure 1E). Responsiveness to jolts differed significantly between resident and visitor clients (Model 5; Analysis of Deviance: Chisq = 15.2, df = 1, p < 0.0001), while growth strategy (Model 5; Analysis of Deviance: Chisq = 5.65, df = 2, p = 0.06), and site (Model 5; Analysis of Deviance: Chisq = 9.34, df = 7, p = 0.23) had no overall effect. None of the interactions resulted in a significant effect (p > 0.1). After model simplification, significances remained, and post hoc analyses indicated that, despite the interaction between growth strategy and client type not being significant, visitor clients were more responsive than resident clients when interacting with slow- (p = 0.004) and average-growing cleaners (p = 0.03), whereas no such difference was observed for fast growers (p = 0.16).

For the PCA (Figure 2A), PC1 described a gradient primarily associated with overall interaction investment (average number of interaction durations and tactile stimulation). Individuals with lower PC1 scores engaged in more frequent and longer interactions and provided more tactile stimulation. PC2 was mainly associated with cheating-related dynamics, contrasting cleaners that elicited more client jolts and responses with those that did not. Fast-, slow-, and average-growing individuals showed substantial overlap in multivariate behavioral space.

**Figure 2:**
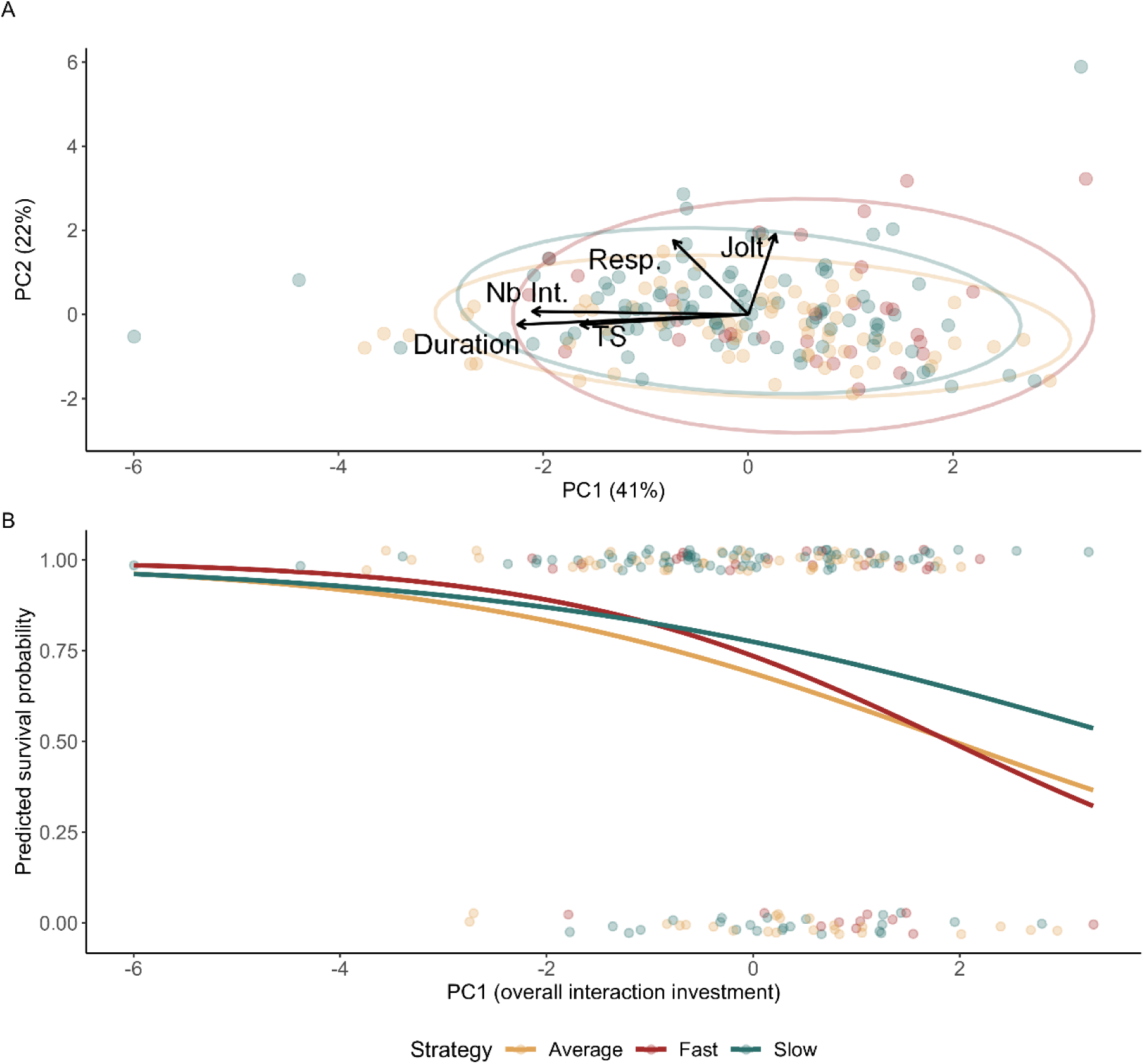
Cleaning behavior and its association with survival probability. *A.* Principal Component Analysis (PCA) of cleaning service characteristics. The PCA summarizes variation in five behavioral variables: average number of interactions (Nb Int.), average interaction duration (Duration), average proportion of interactions with tactile stimulation (TS), average jolt rate (Jolt), and average proportion of jolts that elicited a client’s negative response (Resp.). Points represent individual cleaner wrasse, colored by growth strategy. Arrows indicate variable loadings, and ellipses represent 95% confidence intervals for each growth strategy. PC1 and PC2 explain 41% and 22% of the total variance, respectively. *B.* Predicted survival probability of the three growth strategies based on levels of PC1. Raw individual observations are shown as jitter, and the overlaid lines represent model predictions (Model 6; Binomial GLMM). Survival data include 190 cleaner ID (fast = 30, slow = 91, average = 69).

Survival probability showed an association with PC1 (Model 6; Analysis of Deviance: Chisq = 5.32, df = 1, p = 0.02, Figure 2B), but not with PC2 (Model 6; Analysis of Deviance: Chisq = 0.02, df = 1, p = 0.88), suggesting that interaction investment rather than cheating patterns predict survival probability. The lower the PC1 score, which translates into greater cleaning investment, the higher the survival (Figure 2B). Growth strategy did not significantly influence survival (Model 6; Analysis of Deviance: Chisq = 0.9, df = 2, p = 0.64), and there was no evidence that the relationship between cleaning behavior and survival differed between growth strategies as neither the interaction with PC1 (Model 6; Analysis of Deviance: Chisq = 0.20, df = 2, p = 0.90) nor PC2 (Model 6; Analysis of Deviance: Chisq = 1.76, df = 2, p = 0.42) were significant. For this analysis, behavioural variables were aggregated across client types (i.e., resident and visitor clients were not analysed separately) to avoid model instability and dispersion issues.

### 2. Cleaner densities have an important role

The proportion of time spent being aggressed by conspecifics was not correlated with cleaner density (Pearson’s correlation: Corr = 0.02, t = 0.43, df = 746, p = 0.67). In contrast, the proportion of time spent interacting with larger individuals was weakly but significantly positively correlated with cleaner density (Pearson’s correlation: Corr = 0.12, t = 3.24, df = 746, p = 0.001), indicating a small increase in interaction time with increasing cleaner density.

Proportions of fast, slow, and average growers in the reef were significantly different (Model 8.1; Analysis of Deviance: Chisq = 16.7600, df = 2, p = 0.0002), and this difference was significantly dependent on cleaner-to-client ratio (Model 8.1; Analysis of Deviance: Chisq = 13.709, df = 2, p = 0.0001). Fast- and average-growing strategies have a tendency to be more abundant at high cleaner-to-client ratios, while the slow-growing strategy follows the opposite trend (Figure 3). Post hoc analyses revealed that this trend was significant for the slow growth category (0.03) but not for the fast (0.37) and the average one (0.75). Cleaner fish reef density was moderately but significantly positively correlated with client density (Model 8.2; Pearson’s correlation: Corr = 0.49, t = 2.62, df = 22, p = 0.02). However, the cleaner-to-client ratio was only strongly correlated with cleaner reef density (Model 8.3; Pearson’s correlation: Corr = 0.78, t = 5.78, df = 22, p < 0.0001), and showed no correlation with client Density (Model 8.4; Pearson’s correlation: Corr = -0.16, t = -0.75, df = 22, p = 0.46),.

**Figure 3:**
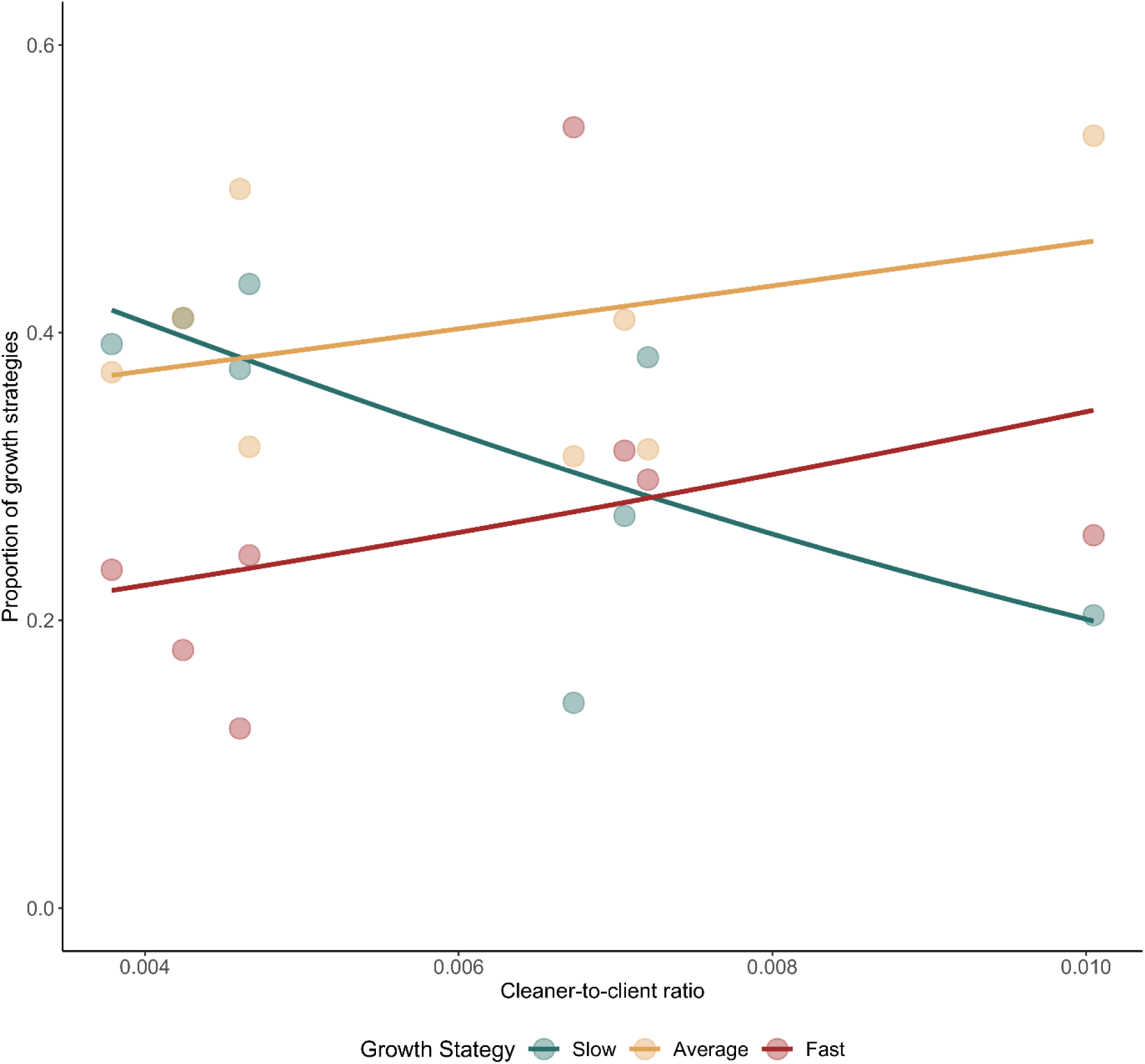
Scatter plot of the relationship between the cleaner-to-client ratio and the proportion of fast, slow, and average growth strategies. The jitter represents raw site-level data (n = 8, strategies = 3), while overlaid lines represent model-estimated means ± standard errors (SE) from Model 8.1. For these two analyses, the general growth curve was used to define the growth strategy, allowing for site differences in cleaner density.

### 3. Lack of evidence for a reproductive trade-off

We tested whether fast-, slow-, and average-growing fish differ in reproductive investment using three measures of reproductive activity: presence of a distended belly, Body-Sigmoid display, and actual spawning.

For the belly measure, there was no effect of growth strategy on the probability of a female showing a distended belly (Model 9; Analysis of Deviance Table: Chisq = 2.25, df = 2, p = 0.3248). Belly was also neither linked to distance to the tide (Model 9; Analysis of Deviance Table: Chisq = 1.03, df = 1, p = 0.3094), nor its quadratic effect (Model 9; Analysis of Deviance Table: Chisq = 1.5127, df = 1, p = 0.2187). Marginal R² for the model was low (0.017), whereas conditional R² was higher (0.292), suggesting that individual variation and lunar phase explained most of the variation in belly occurrence.

For the Body-Sigmoid display, growth strategy again had no effect (Model 10; Analysis of Deviance Table: Chisq = 0.2067, df = 2, p = 0.90182), whereas the quadratic effect of distance to high tide resulted significant (Model 10; Analysis of Deviance Table: Chisq = 6.1194, df = 1, p = 0.01337), with peak display probability occurring roughly 2 hours after high tide. The marginal R² of this model was slightly higher than for belly alone (0.085), as well as the conditional R² (0.416).

For the spawning, growth strategy had no effect (Model 11; Analysis of Deviance Table: Chisq = 1.1813, df = 2, p = 0.55397), whereas the quadratic effect of distance to high tide resulted significant (Model 11; Analysis of Deviance Table: Chisq = 15.4901, df = 1, p <0.0001), with peak display probability occurring roughly 30 minutes after high tide. The marginal R² (0.922) and the conditional R² (0.922) of this model were higher than those of the previous models.

### 4. Social control plays an important role

The proportion of time spent in negative interactions with conspecifics does not seem to be affected by growth strategy (Model 12; Analysis of Deviance: Chisq = 5.0034, df = 2, p = 0.08194; Figure 4A), but by type of aggression alone (Model 12; Analysis of Deviance: Chisq = 25.4417, df = 1, p < 0.0001). However, there was a significant interaction between the effect of growth strategy and aggression type (directed to subordinates or received by dominants) on the proportion of time spent in negative interaction (Model 12; Analysis of Deviance: Chisq = 34.69, df = 2, p < 0.0001). Post hoc analyses showed that slow growers received 79% and 55% more aggression than average and fast growers, respectively. However, only the comparison between slow and average growers was significant (p-value = 0.002).

**Figure 4:**
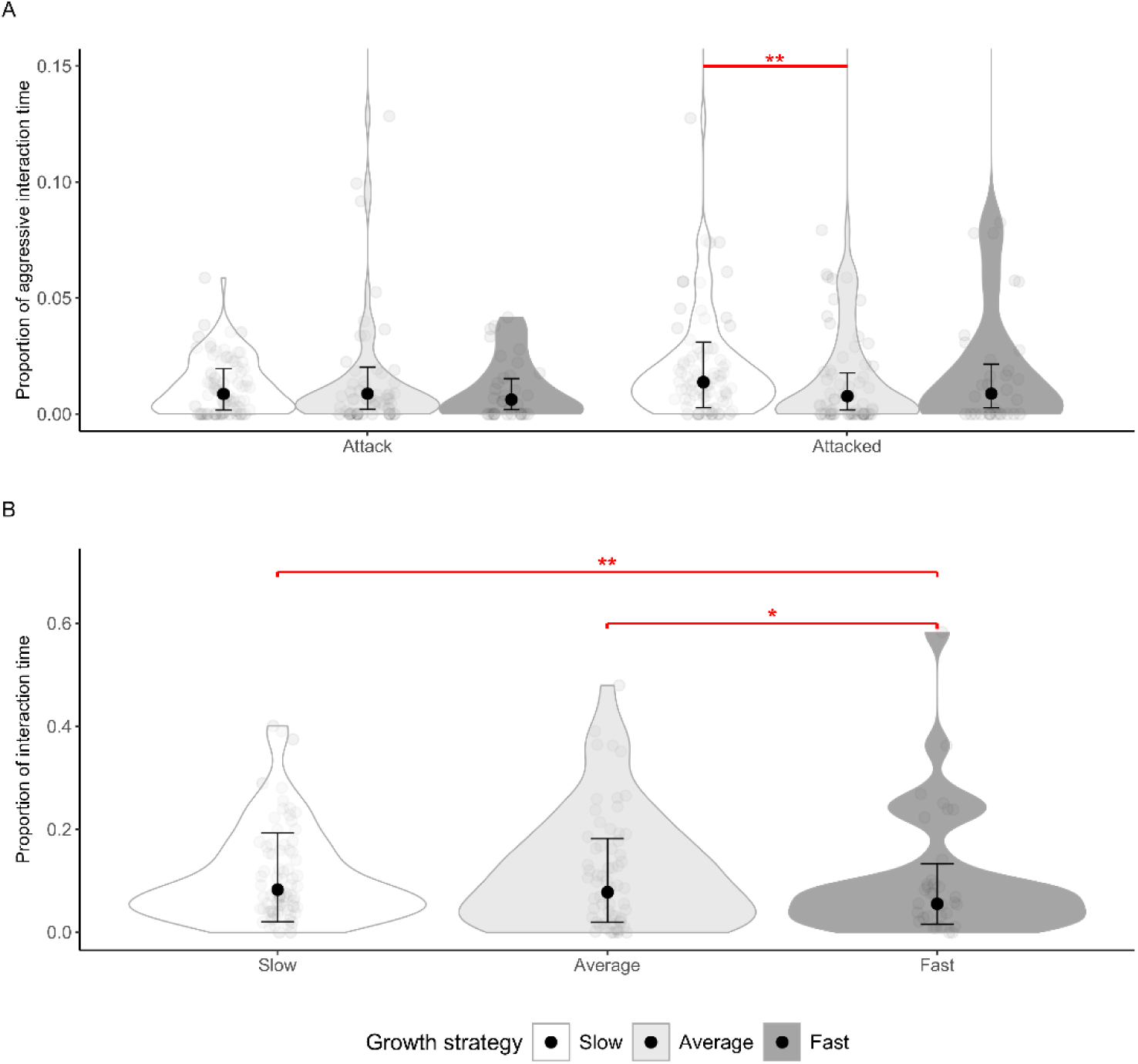
Intraspecific patterns and growth strategy. For each level of growth strategy, these graphs show: *A.* Proportion of time spent in negative interactions by aggression type (attacked by dominants or attack subordinates; Model 12; Gaussian LMM); *B.* Proportion of time spent with larger individuals irrespective of the nature of interaction (Model 13; Gaussian LMM). In both graphs, distribution is shown as violins, the jitter represents raw individual-level data, and overlaid points and error bars represent model-estimated means ± standard errors (SE). Here, local population growth curves were used to define the growth strategy. The line connecting the violins with an asterisk indicates a significant difference between the two groups involved (* *p* < 0.05; ** p < 0.01; *** p < 0.0001). Intraspecific data include 746 videos (fast = 102, slow = 382, average = 262) for 177 cleaner ID (fast = 33, slow = 83, average = 61).

The proportion of general time spent with larger individuals was, however, strongly dependent on growth strategy (Model 13; Analysis of deviance: Chisq = 11.45, df = 2, p = 0.003; Figure 4B). Post hoc analyses revealed that fast growers spent significantly less time with larger individuals than both slow (p = 0.003; 49.3% less) and average growers (p = 0.02; 40% less).

## Discussion

We had asked how foraging strategies, reproductive investment, and hierarchical interactions combine to explain fast and slow growth in cleaner fish *Labroides dimidiatus*, a species well known for the strategic sophistication shown in cleaning interactions with clients. Contrary to our predictions, variation in parameters measuring cleaning behavior did not explain variation in growth patterns, nor did indicators of reproductive activity. According to evidence provided in the study, the degree of social control by dominant harem members emerged as the only factor affecting growth patterns. Notably, slower-growing individuals spent more time in the presence of dominants, despite experiencing similar levels of aggression. This indicates that the mere presence of dominant individuals, rather than active aggression, may be sufficient to regulate subordinate growth. None of our measured parameters interacted with growth patterns to predict mortality risks. Below, we discuss each aspect in more detail.

### Cleaning patterns and densities

Despite our predictions, fast-growing individuals did not appear to obtain more energy because of superior cleaning performance. Instead, two results suggest the opposite, i.e. that slow-growing individuals may obtain more food from cleaning interactions. In this context, the observation that individuals of intermediate growth show largely intermediate results strengthens the view that results are not due to one growth category doing something distinctively different. First, slow growers spent overall more time in cleaning interactions. Time spent cleaning is at least in part affected by clients’ decisions about choosing a cleaner, how long to stay, and how to respond to cleaners’ cheating ^44–49,51^. Second, the observation that, across study demes, individuals at sites with lower competition among cleaners for access to clients (i.e., a lower cleaner-to-client ratio) tended to grow more slowly is consistent with the general conclusion that food intake does not predict growth patterns, thereby contradicting previous hypotheses ^23^. However, as shown in previous research ^54^, cleaner and client densities are positively correlated. This correlation could support the working hypothesis that high fish densities provide a greater abundance of ectoparasites ^34,84^, thereby rendering foraging during cleaning interactions more efficient. Consequently, although fast-growing individuals do not appear to gain more energy through longer or more frequent interactions, they may potentially be more efficient cleaners and thus achieve similar overall energy intake. However, because our data do not allow us to quantify feeding rates in terms of bite rate or foraging efficiency, this remains an open question.

### Lack of reproductive and survival trade-offs

As we did not find evidence that fast growth is due to higher food intake, an alternative explanation could be that fast and slow-growing individuals manage trade-offs differently. However, our data do not provide evidence for this. Our three measures of spawning activity did not yield any evidence that slow growers invest more into current reproduction. It is possible that our proxies merely reflect reproductive readiness rather than actual reproductive investment ^85^. We currently cannot exclude that females following different growth strategies differ in the quantity of eggs produced or in gonadal allocation while maintaining similar levels of spawning behavior. Moreover, reproductive behavior may sometimes serve social functions independent of immediate reproductive investment. In sequential hermaphrodites, the behavioral expression of reproductive roles can precede or occur independently of physiological changes in gonadal function ^12,19^. For example, in some protogynous reef fishes, including the cleaner wrasse, the largest female rapidly adopts male-typical spawning behavior following the disappearance of the dominant male, even while still undergoing gonadal transition ^12,19^. Such decoupling suggests that reproductive behavior may partly function to maintain social position rather than directly reflect reproductive investment ^12,19^. In our system, females may similarly maintain spawning behavior toward the dominant male regardless of their current reproductive investment, either to avoid social conflict or to maintain their position within the reproductive hierarchy. We therefore acknowledge that, ideally, we would have obtained egg count data from spawning events as well (as previously measured by Kuwamura (1981) ^77^. However, the presence of a large belly as an indicator of egg mass did not yield evidence of systematic differences either.

Also, it does not appear that fast growth causes major health risks that would lead to increased mortality, adding to the trade-offs we lack evidence for. As we do not have any physiological measures in a long-term field observation project, it remains an open question whether more subtle physiological costs of fast growth would be identifiable.

### The role of intraspecific interactions

The main emerging explanation for variation in growth rates is that slow growers are constrained by stronger intraspecific control. Interestingly, the mere presence of a dominant individual, rather than active aggression, seems sufficient to regulate subordinate growth. This result aligns with previous findings on sex change in this species at Lizard Island, for which the combined effect of reduced interaction with the harem male and the next largest female would lead to the early sex change of a subordinate ^23^.

This pattern is consistent with predictions from size-based hierarchy theory, in which individuals selectively control those directly below them in rank to maintain a competitive size advantage and reproductive position. In such systems, subordinates are expected to strategically adjust their growth trajectories in response to social pressure ^27,28,30,68–73,86^, thereby reducing the risk of escalated conflict or eviction ^27,28,86–90^. Consistent with this interpretation, fast-growing individuals in our study appeared to experience reduced social control, allowing them to maintain higher growth rates despite comparable ecological conditions.

Failure to control fast-growing individuals may result from multiple mechanisms. At Lizard Island, the cleaner wrasse exhibits two types of social structures: branching and linear. Branching systems are characterized by a larger number of females compared to linear systems ^23^, distributed across several groups (branches) that occupy non-overlapping territories. This spatial separation allows for the coexistence of codominant females, individuals of similar size that avoid interacting with one another. In contrast, linear systems consist of a single group of females of different sizes occupying overlapping territories ^18^.

In branching systems, a codominant female may experience rapid growth due to reduced female-female control and decreased attention from the male, who must divide his time among multiple branches and individuals ^23^. Alternatively, within a branch or in linear systems, a fast-growing individual may occupy a core area located farther from dominant individuals, making it more difficult for them to monitor and regulate her growth. A further possibility is that rapid growth is tolerated when there is a large size gap between the fast-growing individual and the next largest competitor. Finally, some dominant individuals may simply fail to exert effective control. Regardless of the mechanism, such failures in social regulation support the idea that social control plays a key role in shaping growth strategies.

Despite the lack of significance with fast growers, slow growers were still the growth strategy most likely to be pursued more aggressively by higher-rankers (79% and 55% more than average and fast growers, respectively). This tendency, together with the documented increase in social control associated with the presence of dominant individuals, could lead to higher levels of stress. Under chronic stress, fish divert resources from growth toward maintenance and survival ^91,92^, and stress-related neuroendocrine mechanisms can directly suppress somatic growth ^93^. This association makes it unlikely that the cleaner fish system operates as cooperatively breeding fish do. In the latter, an experiment in which both dominant and subordinate individuals were fed ad libitum, subordinates did not increase in length but instead accumulated energy reserves and became heavier ^29,30^. We do not expect slow-growing cleaners to be heavier than fast growers, because the mechanism of stress induction could prevent this.

Regarding between-group intraspecific interactions, because the cleaner-to-client ratio was strongly correlated with cleaner density, our data suggest that sex change is not only a driver of fast growth within a deme but also between demes. Individuals from neighboring harems could indeed represent competitors for access to sex change opportunities ^20,23^.

### Limitation of the study

Like many life history projects, our methodology relied entirely on non-invasive techniques to preserve the study populations and their natural social dynamics. Consequently, we were unable to use otolith extraction to collect true chronological age data for our focal fish. Instead, we relied on model-based estimates of individual ages and deme-specific growth curves constructed from regular size measurements. Furthermore, we can only provide correlational measures of life-history components: cleaning duration and client jolts as indicators of energy gains, and presence/aggression by dominants as measures of exerted social control. With respect to reproductive investment, we had to rely on visual indicators, such as the presence of a large belly, to estimate egg mass and behavioural proxies of spawning activity. In an ideal world, we would have measured caloric intake, body condition, fat reserves, stress hormones, immune parameters, and the number of eggs produced for each spawning event.

## Conclusion

Together, our results point to two main conclusions. First, the potential for rapid growth in slow-growing individuals appears to be socially constrained by intraspecific interactions. Second, fast-growing individuals do not seem to acquire the energy required for accelerated growth through increased cleaning effort or higher service quality. Instead, differences in growth are more likely to arise from variation in how energy is allocated, including potential trade-offs between growth, storage, maintenance, and reproduction. Future work directly measuring body condition, fat reserves, and gonadal energy content would help determine whether differences in physiological allocation underlie the divergence in growth strategies observed here. Integrating these physiological measures with behavioral and social data could provide a more complete understanding of how energy intake, social environment, and strategic allocation interact to shape life history trajectories in sequentially hermaphroditic reef fishes. Beyond physiological research, our results raise a major challenge for cognitive research on cleaner fish: given the documented variation in forebrain size and cell numbers and associated strategic sophistication of cleaners, what are the exact benefits of being smart in a cleaner’s daily life?

## Author contributions and declaration

Conceptualization: LP, RB. Data curation: LP. Formal analysis: LP. Funding acquisition: RB. Investigation: LP. Methodology: LP, RB. Project administration: LP, RB. Resources: LP, RB. Software: LP. Supervision: RB. Validation: LP, RB. Visualization: LP. Writing original draft: LP. Writing, review, and editing: LP, RB. All authors have read and agreed to the submitted version of the manuscript.

## Acknowledgements

We thank the staff of the Lizard Island Research Station in Australia, L. Vail, A. Hoggett, R. Car, and A. Davie, for their exceptional support, for fostering an outstanding research environment, and for their continuous warm welcome. We are grateful to Dr. F. Cortesi and the University of Queensland for hosting, and to Dr. A. Green for training in transect data collection. We also thank A. Viglino, M. Amann, S. Deventer, E. Pattenden, S. Lévy, C. Lebet, and D. Berger for their invaluable assistance in the field, and Dr. Radu Slobodeanu for statistical guidance. A special thanks to the inspiring young researchers we met on the island, whose enthusiasm and thoughtful conversations made the work environment both stimulating and welcoming. This work was supported by the Swiss National Science Foundation (grant 310030_192673/1 to RB).

## Supplementary information

Document S1: this document includes methodological details (Supplement S1-S3 with Figure S1-S4, and Tables S1-S4). It also includes statistical model details (Supplement S4-S15 with Figure S5-S18, and Tables S5-S33)

