## Supplementary Materials for "Social control, not service quality, explains fast growth in the cleaner wrasse *Labroides dimidiatus*"

#### Supplement S1: Study sites and populations

##### Study sites

Fieldwork was conducted at Lizard Island (Great Barrier Reef, Australia) as part of an ongoing long-term monitoring program initiated in July 2022. The present study focuses exclusively on a continuous 11-month period spanning July 2022 to June 2023, during which data were collected year-round.

Eight reef sites were selected based on different densities of cleaner wrasse and client fishes (Fig. S1). Sampling effort was distributed across all months within the study period, capturing both austral winter and summer conditions. To simplify, we defined winter as June to November and summer as December to May. At each of our sites, we created a population of individually identifiable cleaner wrasses by catching and tagging them with visual implant elastomers (VIEs).

**Figure S1: Field observation sites on the coral reefs surrounding Lizard Island, Australia.**

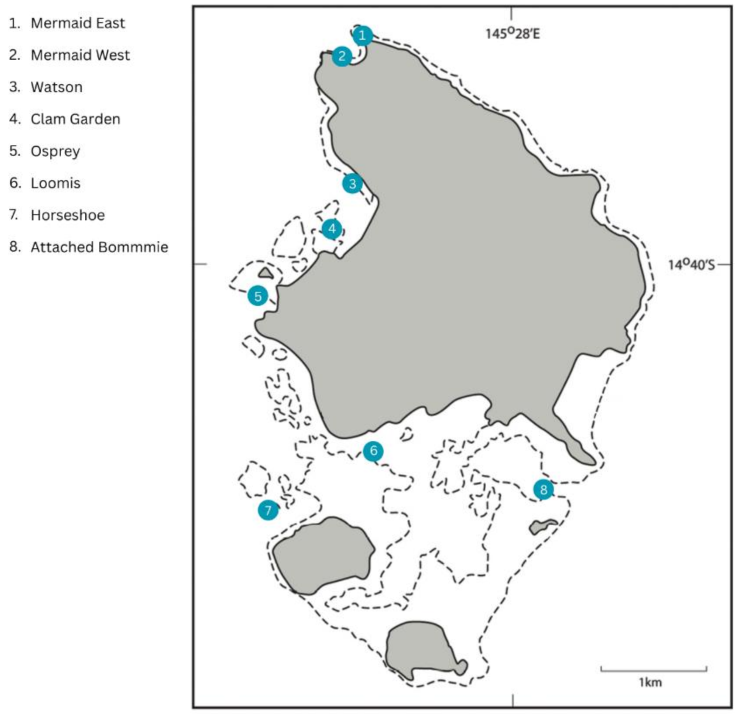

#### Individual identification and tagging protocol.

Cleaner wrasse were captured while SCUBA diving using hand nets (10 x 15 cm) and barrier nets (4.7 y 1.8m or 1 x 1.2m). Handling time was minimized: from capture to release, individuals were restrained in a hand net for less than two minutes. All fish resumed normal behavior immediately after release, and no non-target individuals were captured

Individual identification primarily relied on VIE tags, a widely used technique for small reef fishes [1]. Tagging was conducted underwater to avoid removal from the reef environment. Adult focal individuals received two elastomer injections placed in clear tissue (light band above the black lateral stripe), with injection sites distributed across four possible body locations (anterior/posterior and left/right; Figure S2). Six colors (red, pink, yellow, green, blue, white) were used, yielding up to 1,296 unique tag combinations, with additional differentiation possible when tag order was considered.

Juvenile individuals were marked with a single injection using high-contrast colors (red, pink, or yellow), allowing 108 unique codes. In cases where individuals could be reliably distinguished based on natural markings, such as pigmentation asymmetries or irregularities in the lateral band, tagging was deemed unnecessary (Figure S3).

A total of 280 adults were tagged, and an additional 95 adults were identified based on markings (Table S1). From November 2022, monitoring was expanded to include approximately 505 juveniles. Of these, 165 reached a total length of 40 mm, at which point they were tagged and incorporated into the present dataset. In total, 540 individuals were tracked throughout the study period, with sample sizes ranging from 47 (Loomis) to 88 (Mermaid East).

The primary researcher participated to all fieldwork, totaling ~1,000 dives and ~1,750 hours of underwater observation. This continuous monitoring and field presence enabled consistent recognition of both tagged and untagged individuals. Continuous monitoring enabled the reliable identification of both tagged and untagged individuals.

#### Tag composition and retention

VIE tags consist of a pigmented elastomer that may be combined with a hardening agent to improve structural stability during growth [2]. In this study, the hardener was not used as it sets rapidly at high water temperatures, causing the whole syringe to solidify and go to waste. Despite this, no tag loss was observed over the study period. Some tag expansion occurred as individuals grew, but visibility remained high.

Tag retention was further verified through opportunistic recaptures of previously marked individuals 12–20 months after initial tagging. All recaptured fish retained clearly visible marks. These observations served exclusively to confirm tag durability and are not included in subsequent analyses. Overall, tag persistence exceeded values previously reported for this species [1].

With the exception of two adjacent sites (Mermaid East and Mermaid West), all study locations consisted of isolated reef patches separated by sand or open water. As cleaner wrasse are strongly site-attached and do not traverse open water, movement among sites was not possible. Tag combinations were therefore reused across sites, except between the two Mermaid locations, where unique combinations were maintained.

**Table S1: Total number of focal cleaner wrasses at the 8 sites.**

This table summarizes he total number of tagged (VIE) and non-tagged (No VIE) for the 8 sites: Mermaid East (ME), Mermaid West (MW), Watson (W), Clam Garden (CG), Osprey (O), Horseshoe (H), Loomis (LU), and Attached Bommie (AB).

| Site | Adult Males | | Adult Females | | Juveniles | Total | Sex changers |
|---|---|---|---|---|---|---|---|
| | VIE | No VIE | VIE | No VIE | VIE | | |
| **ME** | 3 | 1 | 45 | 5 | 34 | 88 | 6 |
| **MW** | 6 | 0 | 29 | 10 | 28 | 73 | 7 |
| **W** | 3 | 3 | 31 | 8 | 19 | 64 | 4 |
| **CG** | 5 | 1 | 31 | 9 | 12 | 58 | 5 |
| **O** | 2 | 3 | 33 | 10 | 19 | 67 | 7 |
| **H** | 6 | 0 | 38 | 17 | 18 | 79 | 4 |
| **LU** | 3 | 1 | 18 | 9 | 16 | 47 | 5 |
| **AB** | 5 | 5 | 22 | 13 | 19 | 64 | 4 |
| | 33 | 14 | 247 | 81 | 165 | 540 | 42 |
| **Total** | 47 | | 328 | | 165 | 540 | 42 |

**Figure S2: Illustration of the four locations for VIE**

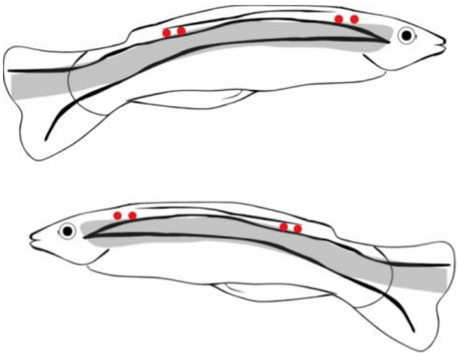

**Figure S3: Examples of two individuals that can be recognized without VIE**

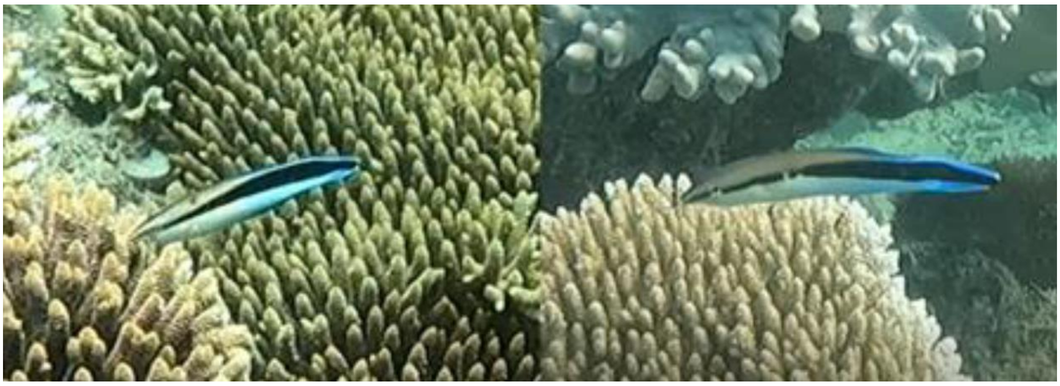

#### Sex identification

Males and females were identified based on behavioral observations in the field, a widely used method in fish behavioral ecology. In this species, males and females exhibit clearly differentiated behaviors, such as territoriality and specific courtship actions, which have been consistently reported in earlier studies and confirmed through physiological measures, including examination of the gonads [3]. Accordingly, the determination of sex in the present study relied on these validated criteria, which included: (i) Flutter-Run, a rapid display in which males swim past females while fluttering their tails and spreading their fins, presenting a lateral profile [3]; (ii) Sexual signaling, where females respond to male courtship with behaviors such as the Body-Sigmoid, a static S-shaped body curve with the belly oriented toward the male, typically indicating readiness to spawn [3]; (iii) Coloration patterns, with females exhibiting distinct sexual colors during courtship that are absent in males [3]; and (iv) Spawning posture, where males assume a dominant position during the later stages of courtship, including straddling the female in an upward spiral and leading the ascent [3]. Beyond these qualitative traits, quantifiable behavioral differences were also noted. Males tolerated the presence of females in close proximity more than females did, especially near feeding areas, and showed higher levels of movement within their territories, frequently visiting females and patrolling territorial boundaries [3]. These behaviors were only used to determine the sex of the focal individuals and were not included as variables in the behavioral analysis described below.

Sex identification was also supported by observations of spawning behavior. Individuals identified as males consistently spawned with smaller partners, whereas females spawned with larger partners. Each spawning event involved only two individuals, ruling out the participation of sneaker males.

#### Supplement S2: Stereo camera sizing error

Following the approach described in Pessina and Bshary (2026a) [4], fish body size was quantified on a monthly basis using a stereo-photogrammetric camera system composed of two GoPro Hero 8 cameras [5]. The paired cameras were mounted on a rigid frame and operated by the primary diver while swimming along the reef, allowing three-dimensional length measurements to be extracted from video recordings. Video footage was processed using EventMeasure software [5], after camera calibration in CAL [5].

Stereo photogrammetry is known to provide substantially greater accuracy than underwater visual estimates [6], with previously reported measurement errors on the order of 1–2 mm [7]. Calibration trials conducted for this study confirmed an overall mean error of ± 1.13mm when measuring a reference bar of known length. Because measuring a moving object is more challenging, the software's accuracy was then investigated by comparing the stereo camera measurements of fish longer than 70cm with manual size measurements taken less than 30 days prior. This method yielded an average error of ±1.81 mm (Figure S4, Table S2). Attempts were made to obtain size measurements using the stereo system within 1 week of the fish's manual post-capture measurement. However, this proved challenging, as the fish required more time to re-acclimate to human presence and often swam too quickly or attempted to escape, making accurate measurements difficult within this short time frame. Nevertheless, the errors associated with our fish measurements did not show a significant increase in variance (variance = 2.324) compared to those observed with the calibration toolbar (variance = 2.264). This suggests that the software's measurement error remains consistent when applied to real fish. The positive shift in the median error for the fish measurements (median = 1.48 mm) reflects the fish's natural growth over the 30 days between the two measurements. This consistency in error variance across both methods indicates that the software performs reliably for measuring wild fish. In cases where sequential measurements indicated an apparent decrease in length, the earlier measurement was retained to conservatively account for measurement uncertainty.

Individuals were typically revisited approximately once per month (median interval = 28 days, mean = 35 days). Changes in body size between consecutive measurements were generally small (median = 1 mm; mean = 2.5 mm). Because the software of the photogrammetric sizing camera records body size as continuous values rather than fixed size classes, measurements did not occur at uniform size intervals among individuals. We therefore used linear interpolation between consecutive measurements to estimate the time required for each fish to reach successive millimeter increments. This procedure reconstructed continuous, evenly spaced size trajectories for each individual, , enabling direct comparisons among fish and facilitating the construction of deme-level size–age growth curves

To derive deme-specific growth rates, we calculated the mean time required for individuals at each site to grow by 1 mm at a given body size. These estimates were then used to infer expected size-age relationships, providing approximate age estimates in the absence of direct age data. Growth trajectories for each deme were subsequently modelled using the von Bertalanffy Growth Function (VBGF), fitted via non-linear least-squares regression with the nlsLM() function from the *minpack.lm* package [8].

**Figure S4: Sizing error using the calibration tool and the focal cleaners**

The following figure shows the size error of the software obtained using sizing focal cleaners ("Cleanerfish") measured 30 days from their manual sizing, and using the objects of known size ("Calibration").

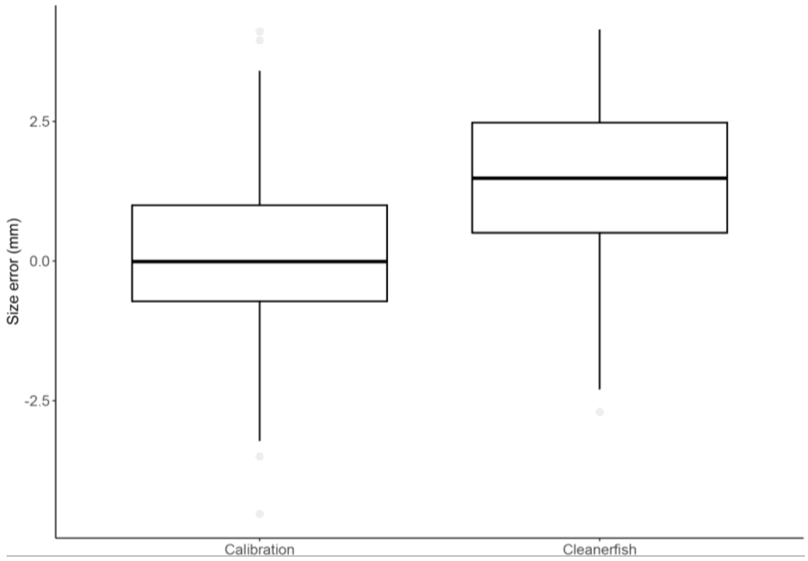

**Table S2: System's errors obtained with the different methods:**

| Method | Mean Error (mm) |
|---|---|
| Small | ± 0.981 |
| Medium | ± 1.08 |
| Large | ± 1.37 |
| General Tool | ± 1.13 |
| Fish | ± 1.81 |

### Supplement S3: Resident and Visitor families

#### Table S3: Residency classification of families

| Family (spp. nb) | Type |
|---|---|
| Acanthuridae (n = 25) | Visitor |
| Apogonidae (n = 17) | Resident |
| Balistidae (n = 7) | Visitor |
| Belonidae (n = 1) | Visitor |
| Blenniidae (n = 12) | Resident |
| Caesionidae (n = 10) | Visitor |
| Caragidae (n =2) | Visitor |
| Carcharhinidae (n = 1) | Visitor |
| Chaetodontidae (n = 24) | Visitor |
| Dasyatidae (n = 1) | Visitor |
| Diodontidae (n = 1) | Visitor |
| Ephippidae (n = 3) | Visitor |
| Gobiidae (n = 7) | Resident |
| Haemulidae (n = 13) | Visitor |
| Hemiraphidae (n = 1) | Visitor |
| Holocentridae (n = 12) | Resident |
| Kyphosidae (n = 2) | Visitor |
| Labridae (n = 50) | Visitor |
| Lethrinidae (n = 9) | Visitor |
| Lutjanidae (n = 12) | Visitor |
| Microdesmidae (n = 2) | Resident |
| Monacanthidae (n = 2) | Resident |
| Mullidae (n = 9) | Visitor |
| Muraenidae (n = 1) | Visitor |
| Nemipteridae (n = 6) | Visitor |
| Ostraciidae (n = 2) | Visitor |
| Pempheridae (n = 1) | Resident |
| Pinguipedidae (n = 4) | Resident |
| Plesiopidae (n = 1) | Resident |
| Pomacanthidae (n = 7) | Visitor |
| Pomacentridae (n = 62) | Resident |
| Priacanthidae (n = 2) | Resident |
| Pseudochromidae (n = 2) | Resident |
| Scaridae (n = 26) | Visitor |
| Scombridae (n = 1) | Visitor |
| Serranidae (n = 25) | Visitor |
| Siganidae (n = 8) | Visitor |
| Sphyraenidae (n = 3) | Visitor |
| Synodontidae (n = 3) | Resident |
| Tetraodontidae (n = 9) | Visitor |
| Zanclidae (n = 1) | Visitor |

#### Supplement S4: Model Details and Assumption Checks

##### Table S4: Glossary for tables S5 and S6

| Term | Definition |
|---|---|
| **Aggression Type** | Attacked by dominants, attacking subordinates. |
| **Belly** | Presence of a distended belly (indication of eggs). |
| **Body Sygmoid** | A behaviour directly implying the presence of eggs. |
| **Chose_Diff** | Distance in time to the nearest high tide (in hours). |
| **Client D.** | Client fish reef density within a deme (per $150m^2$). |
| **Cleaner D.** | Adult cleaner fish reef density within a deme (per $150m^2$). |
| **Duration** | Individual average time spent cleaning during a 20-minute video. |
| **General** | Average proportion of time individuals spend in presence of dominant individuals during a 20 minute video. |
| **ID** | Cleaner ID used as a random factor to control for repeated measures. |
| **Jolt** | Individual average jolt rate. |
| **Moon** | Moon phase. |
| **Number** | Individual average number of cleaning interactions. |
| **Response** | Individual average proportion of jolts that were wither punished or led to termination of the interaction. |
| **Site** | Study deme from which the fish is from. |
| **Strategy** | Growth strategy (fast, slow, or average grower) deriving form deme-specific growth curves. |
| **Strategy g.** | Growth strategy (fast, slow, or average grower) deriving form population-wide growth curve. |
| **Survival** | Binary : alive (1) or dead (o) at the end of the study period. |
| **Type** | Resident or Visitor client. |
| **TS** | Individual average percentage of interactions with tactile stimulation. |

**Table S5: Model Description**

| Model | Description |
|---|---|
| 1 | Investigates differences in average cleaning duration between growth strategies. |
| 2 | Investigates differences in average number of cleaning interactions between growth strategies. |
| 3 | Investigates differences in average jolt rate duration between growth strategies. |
| 4 | Investigates differences in average percentage of cleaning interactions with tactile stimulation duration between growth strategies. |
| 5 | Investigates differences in average client responsiveness between growth strategies. |
| 6 | Investigates survival of fast, slow, and average growers in relation to two principal components formed by our cleaning service variables. |
| 7.1 | Investigates a correlation between cleaner density and the proportion of time spent with dominant individuals. |
| 7.2 | Investigates a correlation between cleaner density and the proportion of aggression received. |
| 8.1 | Investigates if proportion of growth strategies is affected by client reef density and cleaner to client ratio. |
| 8.2 | Investigates a correlations between cleaner density and client density. |
| 8.3 | Investigates a correlation between cleaner density and cleaner-to-client ratio. |
| 8.4 | Investigates a correlation between client density and cleaner-to-client ratio. |
| 9 | Investigates frequency of distended belly (proxy of spawning activity) in fast, slow, and average growers. |
| 10 | Investigates frequency of Body sigmoid (proxy of spawning activity) in fast, slow, and average growers. |
| 11 | Investigates frequency of spawning in fast, slow, and average growers. |
| 12 | Investigates if the average proportion of aggression directed and received is affected by cleaner fish reef density for both growth strategies. |
| 13 | Investigates differences in the average proportion of time spent with larger individuals between growth strategies. |

#### Table S6: Model Summary

| Mode | Type | Family(link) | Formula | Nb of Video(nb of ID) |
|---|---|---|---|---|
| 1 | LMM | Gaussian | log(Duration+3.05)~Strategy+Type+Site+Strategy:Type+ Strategy:Site+Type:Strategy+(1\|ID), dispformula = ~Site | Fast =120(34), Average = 323(67), Slow = 471(93) |
| Simplified to: | | | log(Duration+3.05)~Strategy+Type+Site +Type:Strategy+(+\|ID), dispformula = ~Site*Type+Strategy | |
| 2 | LMM | Gaussian | log(Number+2.9)~Strategy+Type+Site+Strategy:Type+Strategy:Site +Type:Site+(1\|ID) | Fast =120(34), Average = 323(67), Slow = 471(93) |
| Simplified to: | | | log(Number+2.9)~Strategy+Type+Site+Type:Site+(1\|ID) | |
| 3 | LMM | Gaussian | log(Jolt+1.61)~Strategy+Type+Site+Strategy:Type+Strategy:Site+Ty pe:Site+(1\|ID) | Fast =120(34), Average = 323(67), Slow = 471(93) |
| Simplified to: | | | log(Jolt+1.61)~Strategy+Type+Site+(1\|ID) | |
| 4 | GLMM | Beta(logit) | TS~Strategy+Type+ Site+ Strategy:Site+Strategy:Type+ Type:Site+(1\|ID), dispformula =~Strategy+Type+Site | Fast =120(34), Average = 323(67), Slow = 471(93) |
| Simplified to: | | | TS~Strategy+Type+ Site+ Type:Site+(1\|ID), dispformula =~Strategy+Type+Site | |
| 5 | GLMM | Tweedie(log) | Response ~ Strategy +Type + Site+ Strategy:Type + Strategy:Site + Type:Site + (1 \| ID) | Fast =120(34), Average = 323(67), Slow = 471(93) |
| Simplified to: | | | Response ~ Strategy *Type + Site + (1 \| ID) | |
| 6 | GLMM | Binomial(logit) | Survival ~(PC1 + PC2)*Strategy + (1\|Site) | Average = 69, Slow =91, Fast = 30 |
| 7.1 | Cor.test | | cor.test(General, Cleaner d.) | n=748 id=177 |
| 7.2 | Cor.test | | cor.test(Attacked, Cleaner d.) | n=748 id=177 |
| 8.1 | GLMM | Binomial(logit) | Prop~Strategy*Ratio, weights = Tot | Sites = 8 |
| 8.2 | Cor.test | | Cor.test(Cleaner d., Client d.) | Sites =8 |
| 8.3 | Cor.test | | Cor.test(Ratio, Cleaner d., ) | Sites =8 |
| 8.4 | Cor.test | | Cor.test(ratio, Client d.,) | Sites =8 |
| 9 | GLMM | Binomial(logit) | Belly ~ Strategy +Chose_Diff + I(Chose_Diff^2) +(1\|Site) +(1 \| ID)+(Chose_Diff \| Moon) | Fast=87(32), Avg = 188(62), Slow= 286(83) |
| 10 | GLMM | Binomial(logit) | Body Sygmoid ~ Strategy +Chose_Diff + I(Chose_Diff^2) +(1\|Site)+(1 \| ID)+(Chose_Diff \| Moon) | Fast=87(32), Avg = 188(62), Slow= 286(83) |
| 11 | GLMM | Binomial(logit) | Spawn ~ Strategy +Chose_Diff + I(Chose_Diff^2) +(1\|Site)+(1\|Moon) | Fast=87(32), Avg = 188(62), Slow= 286(83) |
| 12 | LMM | Gaussian | Log(Attacked) ~Strategy * Aggression Type+(1\|Site/ID), weights = number videos. | Fast=102( 33), Slow = 382(83), Avg = 262(61) |
| 13 | LMM | Gaussian | Log(General)~Strategy+(1\|Site), weights = number of video | Fast=102( 33), Slow = 382(83), Avg = 262(61) |

#### Supplement S5: Details for Model 1

##### Figure S5: Diagnostic Plots for Model 1

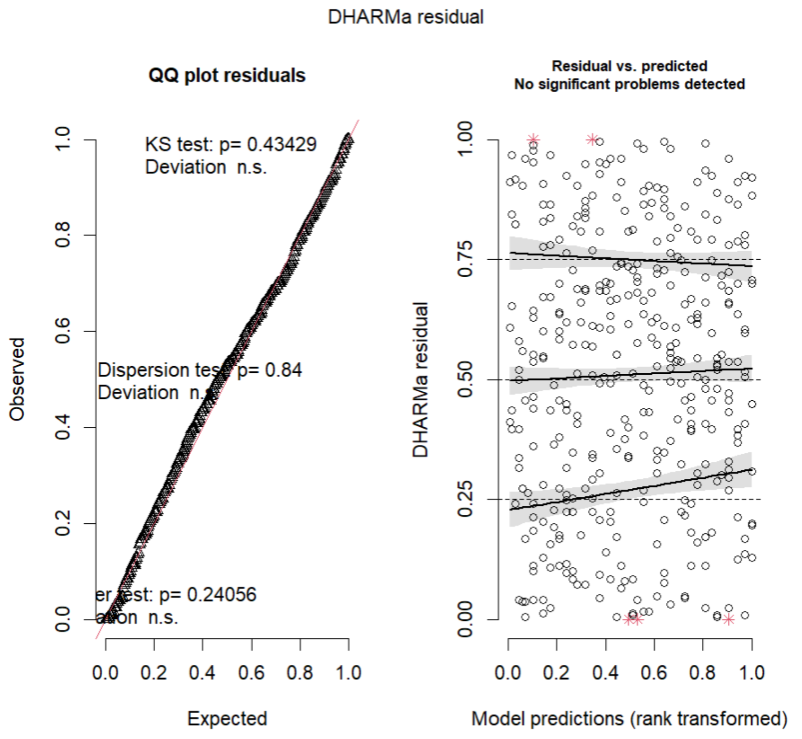

##### Table S7: Analyss of deviance: Type II Wald Chi-square Tests – Model 1

| Term | Chisq | Df | Pr(>Chisq) |
|---|---|---|---|
| Strategy | 15.1590 | 2 | 0.0005108** |
| Type | 214.5500 | 1 | <2.2e-16*** |
| Site | 21.4119 | 7 | 0.0032060** |
| Strategy:Site | 16.4789 | 14 | 0.2850118 |
| Strategy:Type | 1.1243 | 2 | 0.5699758 |
| Type:Site | 27.8120 | 7 | 0.0002378*** |

##### Table S8: First Emmeans Contrast for Model 1

| **Contrast Resident-Visitor** | | | | | |
|---|---|---|---|---|---|
| **Contrast** | **estimate** | **SE** | **df** | **t.ratio** | **p-value** |
| CG | -1.080 | 0.105 | 347 | -10.3 | <.0001*** |
| H | -0.321 | 0.181 | | -1.772 | 0.0772 |
| LU | -0.350 | 0.296 | | -1.182 | 0.2379 |
| AB | -1.141 | 0.238 | | -4.786 | <.0001*** |
| ME | -1.376 | 0.189 | | -7.268 | <.0001*** |
| MW | -1.139 | 0.218 | | -5.234 | <.0001*** |
| O | -0.712 | 0.179 | | -3.977 | 0.0001*** |
| W | -0.483 | 0.206 | | -2.339 | 0.0199* |

Contrasts are still on the log(mu + 3.5) scale.

**Table S9: Second Emmeans Contrast for Model 1**

| **Contrast Resident-Visitor** | | | | | |
|---|---|---|---|---|---|
| **Contrast** | **estimate** | **SE** | **df** | **t.ratio** | **p-value** |
| Slow-Average | 0.161 | 0.0912 | 347 | 1.771 | 0.1810 |
| Slow-Fast | 0.401 | 0.1130 | 347 | 3.538 | 0.0013** |
| Average-Fast | 0.239 | 0.1240 | 347 | 1.933 | 0.1310 |

Results are averaged over the levels of: Type, Site

Note: contrasts are still on the log(mu + 3.05555555555556) scale.

**Figure S6: Effect of site on average cleaning time.**

For each level of client type (Resident or Visitor) and growth strategies, this graph shows the distribution of average cleaning duration for each site and each client type as violin plots. Jittered points represent raw individual-level data. The overlaid points and error bars indicate model-estimated means ± standard errors (SE; Model 1; Gaussian LMM). The asterisk indicates a significant difference between the two groups involved (* $p < 0.05$; ** $p < 0.01$; *** $p < 0.0001$). Cleaning data includes 914 videos (fast = 120, slow = 471, average = 323) for 194 cleaner ID (fast = 34, slow = 93, average = 67).

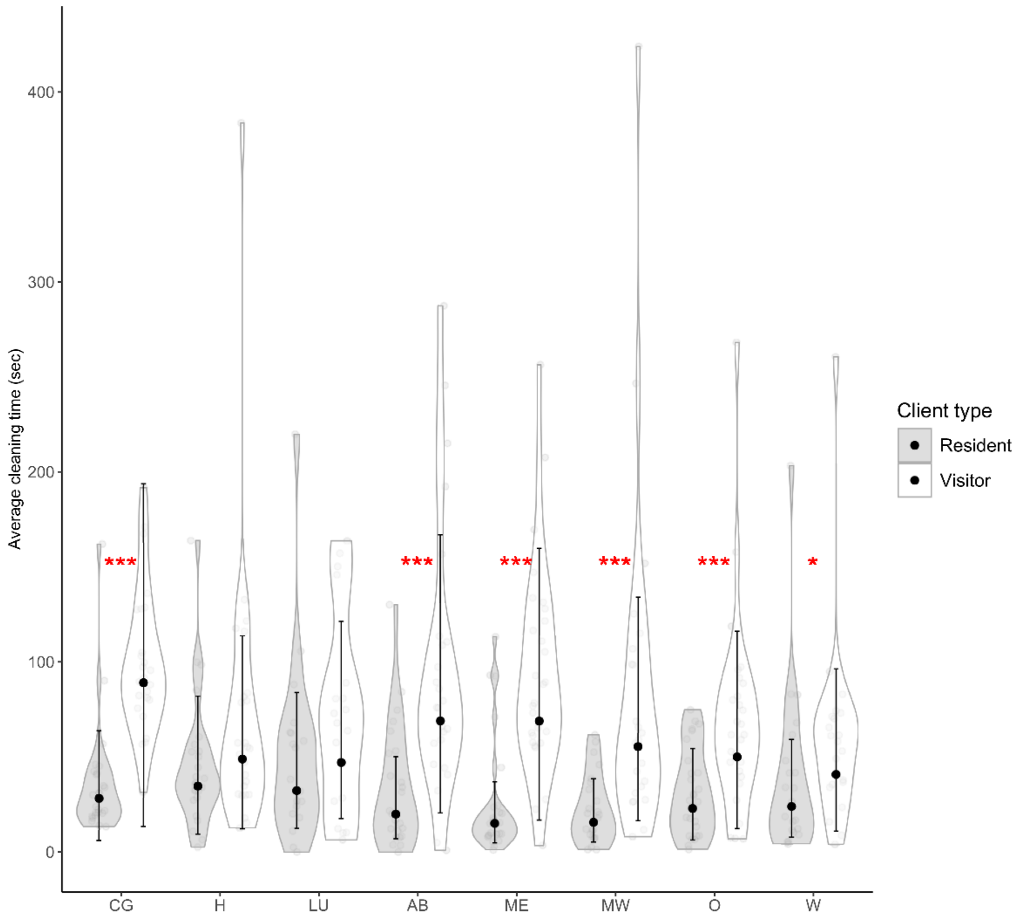

#### Supplement S6: Details for Model 2

##### Figure S7: Diagnostic Plots for Model 2

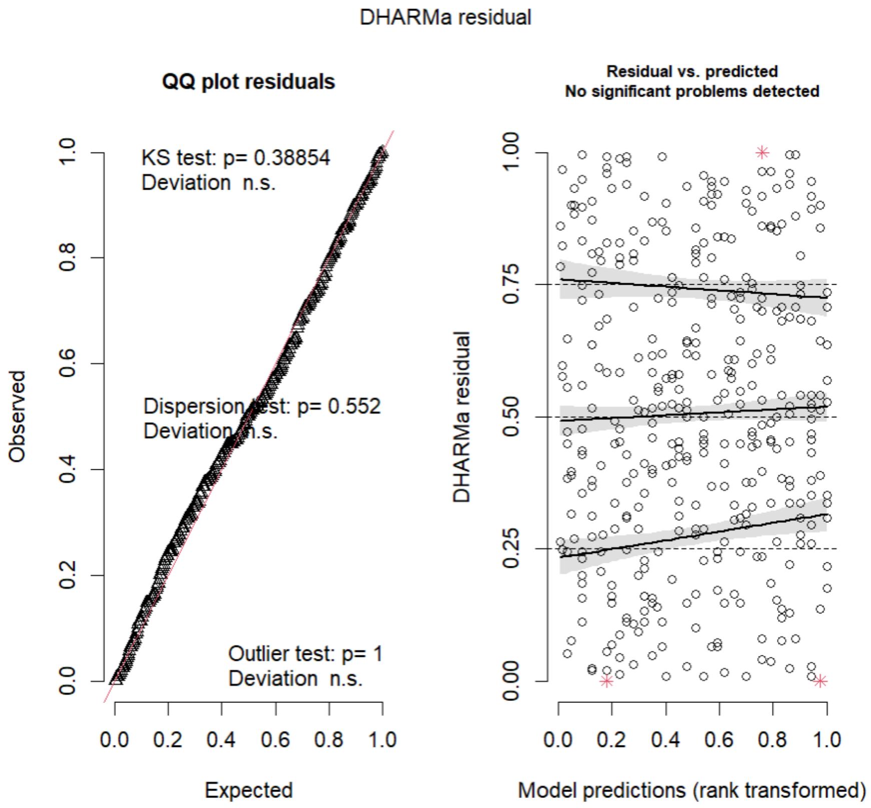

##### Table S10: Analyss of deviance - Type II Wald Chi-square Tests for Model 2

| Term | Chisq | Df | Pr(>Chisq) |
|---|---|---|---|
| Strategy | 8.5892 | 2 | 0.01364 * |
| Type | 18.8748 | 1 | 1.396e-05*** |
| Site | 14.8732 | 7 | 0.03776* |
| Strategy:Type | 2.2924 | 2 | 0.31784 |
| Strategy:Site | 13.0641 | 14 | 0.52184 |
| Type:Site | 67.0498 | 7 | 5.818e-12 *** |

**Table S11: First Emmeans Contrast for Model 2**

| Contrast Resident-Visitor | | | | | |
|---|---|---|---|---|---|
| **Site** | **estimate** | **SE** | **Df** | **t.ratio** | **p-value** |
| CG | -0.330 | 0.111 | 182 | -2.978 | 0.0033** |
| H | 0.222 | 0.111 | 182 | 2.006 | 0.0463* |
| LU | 0.159 | 0.130 | 186 | 1.224 | 0.2224 |
| AB | -0.113 | 0.113 | 185 | -1.002 | 0.3178 |
| ME | -0.685 | 0.106 | 185 | -6.456 | <.0001*** |
| MW | -0.559 | 0.116 | 182 | -4.840 | <.0001*** |
| O | -0.111 | 0.106 | 185 | -1.048 | 0.2958 |
| W | 0.193 | 0.121 | 182 | 1.599 | 0.1116 |

Contrasts are still on the log(mu + 2.9) scale.

**Figure S8: Effect of site on average number of interactions cleaning time.**

For each level of client type (Resident or Visitor) and growth strategies, this graph shows the distribution of average number of cleaning interactions for each site and each client type as violin plots. Jittered points represent raw individual-level data. The overlaid points and error bars indicate model-estimated means ± standard errors (Model 2: Gaussian LMM). The asterisk indicates a significant difference between the two groups involved (* $p < 0.05$; ** $p < 0.01$; *** $p < 0.0001$). Cleaning data includes 914 videos (fast = 120, slow = 471, average = 323) for 194 cleaner ID (fast = 34, slow = 93, average = 67).

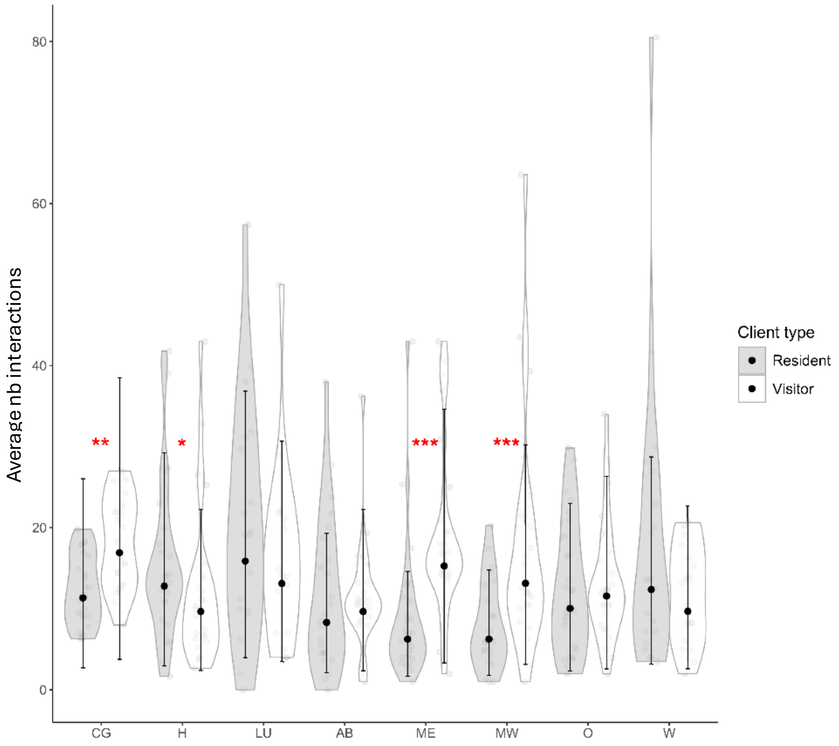

##### Table S12: Second Emmeans Contrast for Model 2

| Contrast | estimate | SE | df | t.ratio | P value |
|---|---|---|---|---|---|
| Average-Fast | 0.133 | 0.0972 | 185 | 1.374 | 0.3567 |
| Average-Slow | -0.127 | 0.0733 | 183 | -1.732 | 0.1961 |
| Fast-Slow | -0.261 | 0.0925 | 185 | -2.816 | 0.0149* |
| contrasts are still on the log(mu + 2.9) scale. | | | | | |

#### Supplement S7: Details for Model 3

##### Figure S9: Diagnostic Plots for Model 3

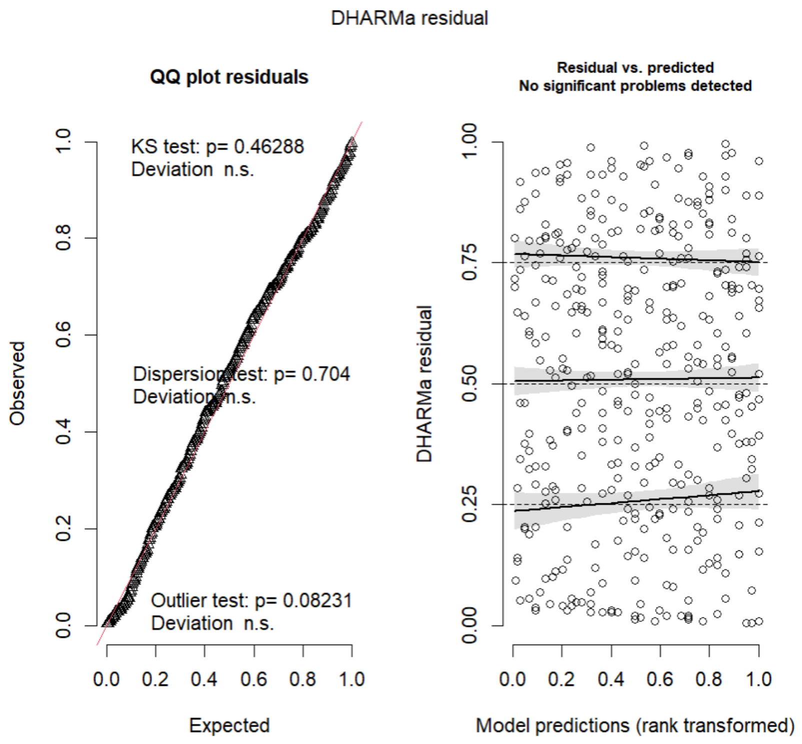

##### Table S13: Analyss of deviance - Type II Wald Chi-square Tests for Model 3

| Term | Chisq | Df | Pr(>Chisq) |
|---|---|---|---|
| Strategy | 13.1738 | 2 | 0.001378** |
| Type | 9.6372 | 1 | 0.001907** |
| Site | 5.1440 | 7 | 0.642395 |
| Strategy:Type | 2.3844 | 2 | 0.303552 |
| Strategy:Site | 18.6959 | 14 | 0.176897 |
| Type:Site | 10.7936 | 7 | 0.147878 |

**Table S14: Firs Emmeans Contrast for Model 3**

| Contrast | estimate | SE | df | t.ratio | p-value |
|---|---|---|---|---|---|
| Slow-Average | 0.299 | 0.108 | 182 | 2.764 | 0.0172* |
| Slow-Fast | 0.421 | 0.137 | 185 | 3.076 | 0.0068* |
| Average_fast | 0.122 | 0.144 | 185 | 0.846 | 0.6746 |
| Contrasts are still on the log(mu + 1.61) scale. | | | | | |

**Table S15: Second Emmeans Contrast for Model 3**

| Contrast | estimate | SE | df | t.ratio | P value |
|---|---|---|---|---|---|
| Resident-Visitor | 0.249 | 0.0806 | 192 | 3.083 | 0.0024** |
| Contrasts are still on the log(mu + 1.61) scale. | | | | | |

#### Supplement S8: Details for Model 4

##### Figure S10: Diagnostic Plots for Model 4

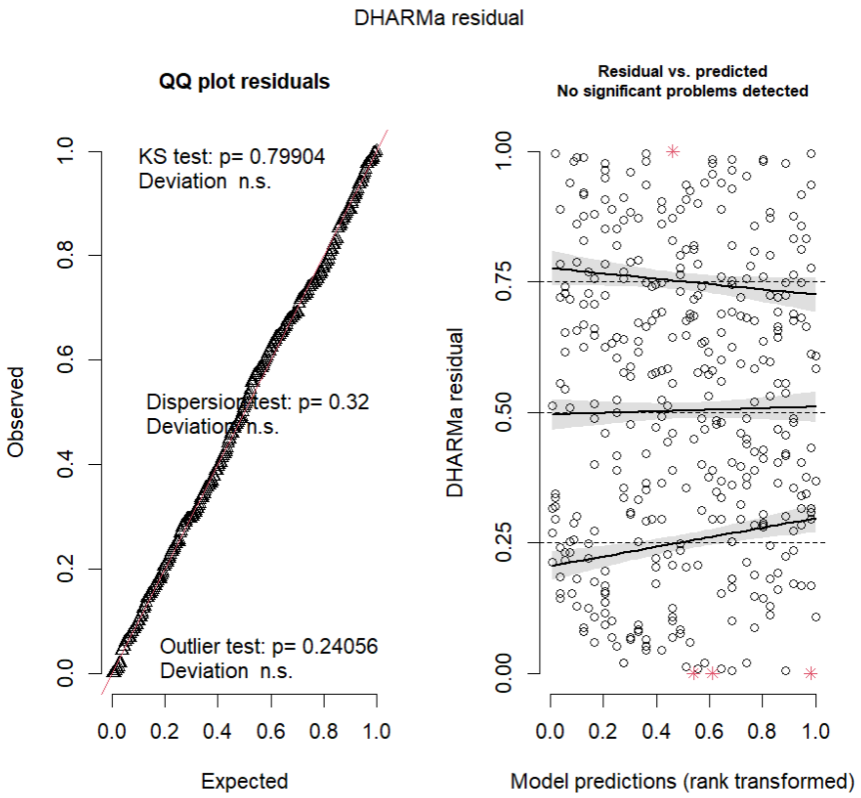

**Table S16: Analyss of deviance - Type II Wald Chi-square Tests for Model 4**

| Term | Chisq | Df | Pr(>Chisq) |
|---|---|---|---|
| Strategy | 3.9821 | 2 | 0.13655 |
| Type | 223.0188 | 1 | <2e-16*** |
| Site | 14.1353 | 7 | 0.04883* |
| Strategy:Site | 16.4819 | 14 | 0.28484 |
| Strategy:Type | 1.2688 | 2 | 0.53025 |
| Type:Site | 6.0246 | 7 | 0.53688 |

**Table S17: First Emmeans Contrast for Model 4**

| Contrast | estimate | SE | df | z.ratio | p-value |
|---|---|---|---|---|---|
| Slow-Average | -0.058 | 0.131 | Inf | -0.442 | 0.8980 |
| Slow-Fast | 0.271 | 0.172 | Inf | 1.575 | 0.2566 |
| Average_fast | 0.329 | 0.183 | Inf | 1.794 | 0.1714 |
| Results are averaged over the levels of : Strategy, Site. | | | | | |
| Results are given on the log odds ratio (not the response) scale. | | | | | |

**Table S18: Second Emmeans Contrast for Model 4**

| Contrast | estimate | SE | df | z.ratio | P value |
|---|---|---|---|---|---|
| Resident-Visitor | -1.47 | 0.101 | Inf | -14.578 | <.0001*** |
| Results are averaged over the levels of : Strategy, Site. | | | | | |
| Results are given on the log odds ratio (not the response) scale. | | | | | |

### Supplement S9: Details for Model 5

#### Figure S11: Diagnostic Plots for Model 5

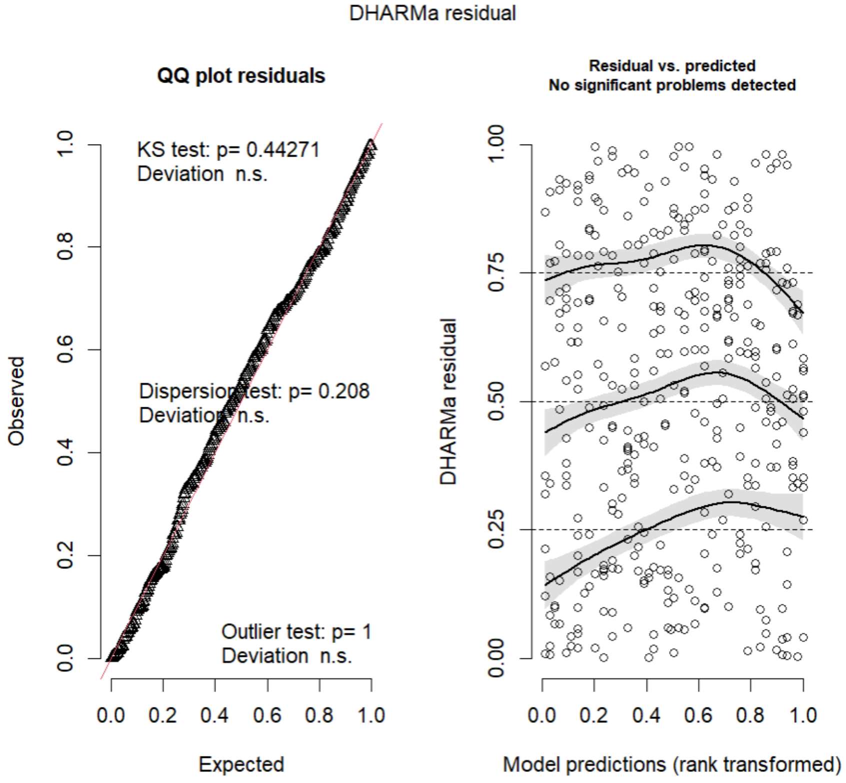

#### Table S19: Analyss of deviance - Type II Wald Chi-square Tests for Model 5

| Term | Chisq | Df | Pr(>Chisq) |
|---|---|---|---|
| Strategy | 5.6484 | 2 | 0.05936 . |
| Type | 15.2009 | 1 | 9.666e-05*** |
| Site | 9.3391 | 7 | 0.22922 |
| Strategy:Site | 13.1226 | 14 | 0.51690 |
| Strategy:Type | 0.2381 | 2 | 0.88775 |
| Type:Site | 9.1693 | 7 | 0.24073 |

**Table S20: First Emmeans Contrast for Model 5**

| Resident/Visitor | estimate | SE | df | z.ratio | P value |
|---|---|---|---|---|---|
| | | Type = Resident | | | |
| Average/Fast | 0.803 | 0.159 | Inf | -1.109 | 0.5086 |
| Average/Slow | 0.785 | 0.121 | Inf | -1.573 | 0.2574 |
| Fast/Slow | 0.978 | 0.180 | Inf | -0.122 | 0.9919 |
| | | Type = Visitor | | | |
| Average/Fast | 0.848 | 0.150 | Inf | -0.932 | 0.6200 |
| Average/Slow | 0.776 | 0.105 | Inf | -1.885 | 0.1430 |
| Fast/Slow | 0.914 | 0.150 | Inf | -0.544 | 0.8495 |

Results are averaged over the levels of: Site
Tests are performed on the log scale

**Table S21: First Emmeans Contrast for Model 5**

| Resident/Visitor | estimate | SE | df | z.ratio | P value |
|---|---|---|---|---|---|
| Strategy = Fast | 0.757 | 0.1490 | Inf | -1.409 | 0.1588 |
| Strategy = Average | 0.717 | 0.1090 | Inf | -2.191 | 0.0285* |
| Strategy = Slow | 0.708 | 0.0837 | Inf | -2.920 | 0.0035** |

Results are averaged over the levels of: Site
Tests are performed on the log scale

#### Supplement S10: Details for Model 6

##### Figure S12: Diagnostic Plots for Model 6

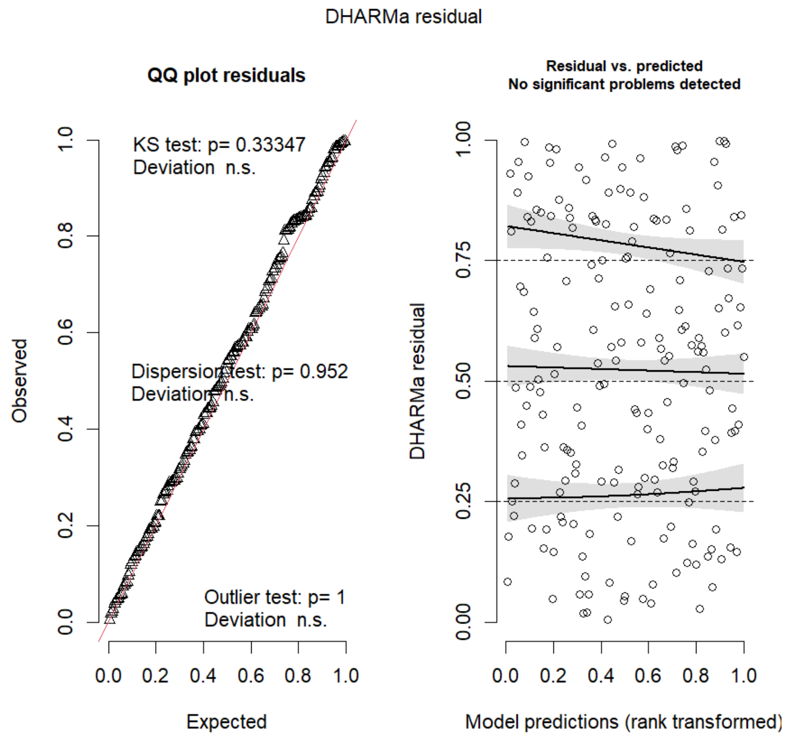

##### Table S22: Analyss of deviance - Type II Wald Chi-square Tests for Model 6

| Term | Chisq | Df | Pr(>Chisq) |
|---|---|---|---|
| PC1 | 5.3176 | 1 | 0.02111* |
| PC2 | 0.0225 | 1 | 0.88077 |
| Strategy | 0.8998 | 2 | 0.63769 |
| PC1:Strategy | 0.2070 | 2 | 0.90170 |
| PC2:Strategy | 1.7580 | 2 | 0.41520 |

#### Supplement S11: Details for Model 7

##### Table S23: Pearson's product-moment correlations for Model 7.1 and 7.2

| Contrast | corr | t | df | p-value |
|---|---|---|---|---|
| 7.1: General/Cleaner D. | 0.1176949 | 3.2371 | 746 | 0.001261** |
| 7.2: Attacked/Cleaner D. | 0.01581289 | 0.43195 | 746 | 0.6659 |

#### Supplement S12: Details for Model 8.1

##### Figure S13: Diagnostic Plots for Model 8.1

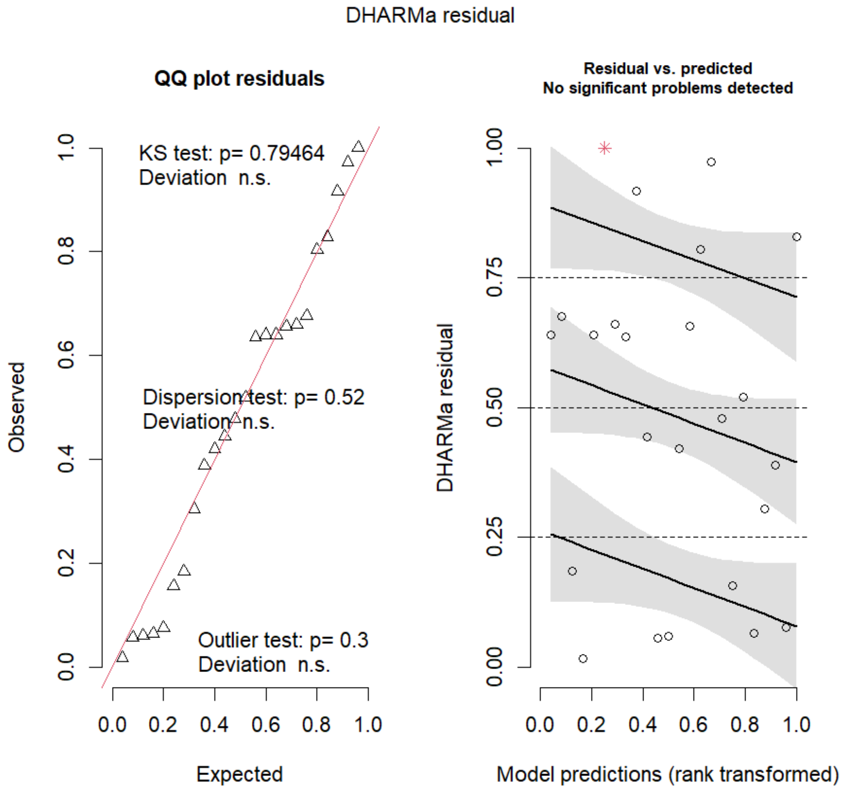

##### Table S24: Analyss of deviance - Type II Wald Chi-square Tests for Model 8.1

| Term | LR.Chisq | Df | Pr(>Chisq) |
|---|---|---|---|
| Strategy general | 16.7600 | 2 | 0.000022294*** |
| Ratio | 0.0.000 | 1 | 0.9999997 |
| Strategy general :Ratio | 13.709 | 2 | 0.00010545*** |

#### Table S25: Emmtrend contrast for Model 8.1

```
$contrasts
Strategy = Slow:
 contrast                                                odds.ratio     SE  df null z.ratio p.value
 Ratio0.00378818627053061 / Ratio0.00535262403580206          1.300 0.1170 Inf    1   2.916  0.0292
 Ratio0.00378818627053061 / Ratio0.006917061801073 5          1.689 0.3040 Inf    1   2.916  0.0292
 Ratio0.00378818627053061 / Ratio0.00848149956634495          2.195 0.5920 Inf    1   2.916  0.0292
 Ratio0.00378818627053061 / Ratio0.010045937331616 4          2.853 1.0300 Inf    1   2.916  0.0292
 Ratio0.00535262403580206 / Ratio0.006917061801073 5          1.300 0.1170 Inf    1   2.916  0.0292
 Ratio0.00535262403580206 / Ratio0.00848149956634495          1.689 0.3040 Inf    1   2.916  0.0292
 Ratio0.00535262403580206 / Ratio0.010045937331616 4          2.195 0.5920 Inf    1   2.916  0.0292
 Ratio0.006917061801073 5 / Ratio0.00848149956634495          1.300 0.1170 Inf    1   2.916  0.0292
 Ratio0.006917061801073 5 / Ratio0.010045937331616 4          1.689 0.3040 Inf    1   2.916  0.0292
 Ratio0.00848149956634495 / Ratio0.010045937331616 4          1.300 0.1170 Inf    1   2.916  0.0292

Strategy = Average:
 contrast                                                odds.ratio     SE  df null z.ratio p.value
 Ratio0.00378818627053061 / Ratio0.00535262403580206          0.908 0.0723 Inf    1  -1.209  0.7463
 Ratio0.00378818627053061 / Ratio0.006917061801073 5          0.825 0.1310 Inf    1  -1.209  0.7463
 Ratio0.00378818627053061 / Ratio0.00848149956634495          0.749 0.1790 Inf    1  -1.209  0.7463
 Ratio0.00378818627053061 / Ratio0.010045937331616 4          0.681 0.2170 Inf    1  -1.209  0.7463
 Ratio0.00535262403580206 / Ratio0.006917061801073 5          0.908 0.0723 Inf    1  -1.209  0.7463
 Ratio0.00535262403580206 / Ratio0.00848149956634495          0.825 0.1310 Inf    1  -1.209  0.7463
 Ratio0.00535262403580206 / Ratio0.010045937331616 4          0.749 0.1790 Inf    1  -1.209  0.7463
 Ratio0.006917061801073 5 / Ratio0.00848149956634495          0.908 0.0723 Inf    1  -1.209  0.7463
 Ratio0.006917061801073 5 / Ratio0.010045937331616 4          0.825 0.1310 Inf    1  -1.209  0.7463
 Ratio0.00848149956634495 / Ratio0.010045937331616 4          0.908 0.0723 Inf    1  -1.209  0.7463

Strategy = Fast:
 contrast                                                odds.ratio     SE  df null z.ratio p.value
 Ratio0.00378818627053061 / Ratio0.00535262403580206          0.855 0.0743 Inf    1  -1.798  0.3747
 Ratio0.00378818627053061 / Ratio0.006917061801073 5          0.732 0.1270 Inf    1  -1.798  0.3747
 Ratio0.00378818627053061 / Ratio0.00848149956634495          0.626 0.1630 Inf    1  -1.798  0.3747
 Ratio0.00378818627053061 / Ratio0.010045937331616 4          0.535 0.1860 Inf    1  -1.798  0.3747
 Ratio0.00535262403580206 / Ratio0.006917061801073 5          0.855 0.0743 Inf    1  -1.798  0.3747
 Ratio0.00535262403580206 / Ratio0.00848149956634495          0.732 0.1270 Inf    1  -1.798  0.3747
 Ratio0.00535262403580206 / Ratio0.010045937331616 4          0.626 0.1630 Inf    1  -1.798  0.3747
 Ratio0.006917061801073 5 / Ratio0.00848149956634495          0.855 0.0743 Inf    1  -1.798  0.3747
 Ratio0.006917061801073 5 / Ratio0.010045937331616 4          0.732 0.1270 Inf    1  -1.798  0.3747
 Ratio0.00848149956634495 / Ratio0.010045937331616 4          0.855 0.0743 Inf    1  -1.798  0.3747

P value adjustment: tukey method for comparing a family of 5 estimates
Tests are performed on the log odds ratio scale
```

#### Table S26: Pearson's product-moment correlations for Model 8.2, 8.3, and 8.4

| Contrast | corr | t | df | p-value |
|---|---|---|---|---|
| Cleaner D./Ratio | 0.7766087 | 5.7821 | 22 | 8.121e-06*** |
| Client C./Ratio | -0.156875 | -0.74503 | 22 | 0.4641 |
| Cleaner D./Client D. | 0.4881177 | 2.6232 | 22 | 0.01553* |

#### Supplement S13: Details for Model 9

##### Figure S14: Diagnostic Plots for Model 9

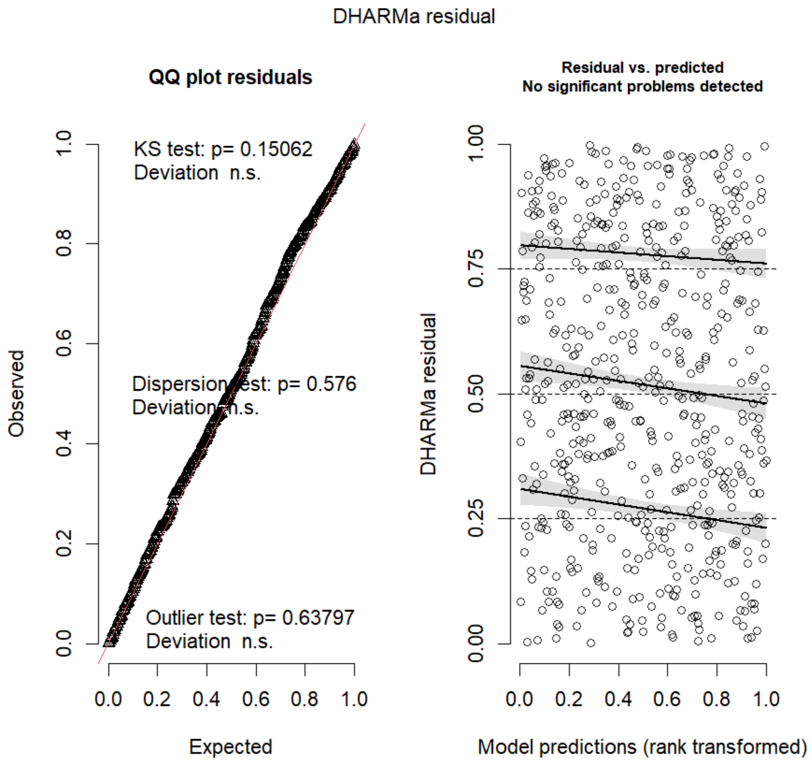

##### Table S27: Analyss of deviance - Type II Wald Chi-square Tests for Model 9

| Term | Chisq | Df | Pr(>Chisq) |
|---|---|---|---|
| Strategy | 2.2389 | 2 | 0.3248 |
| Chose_Diff | 1.0332 | 1 | 0.3094 |
| I(Chose_Diff^2) | 1.5127 | 1 | 0.2187 |

#### Supplement S14: Details for Model 10

##### Figure S15: Diagnostic Plots for Model 10

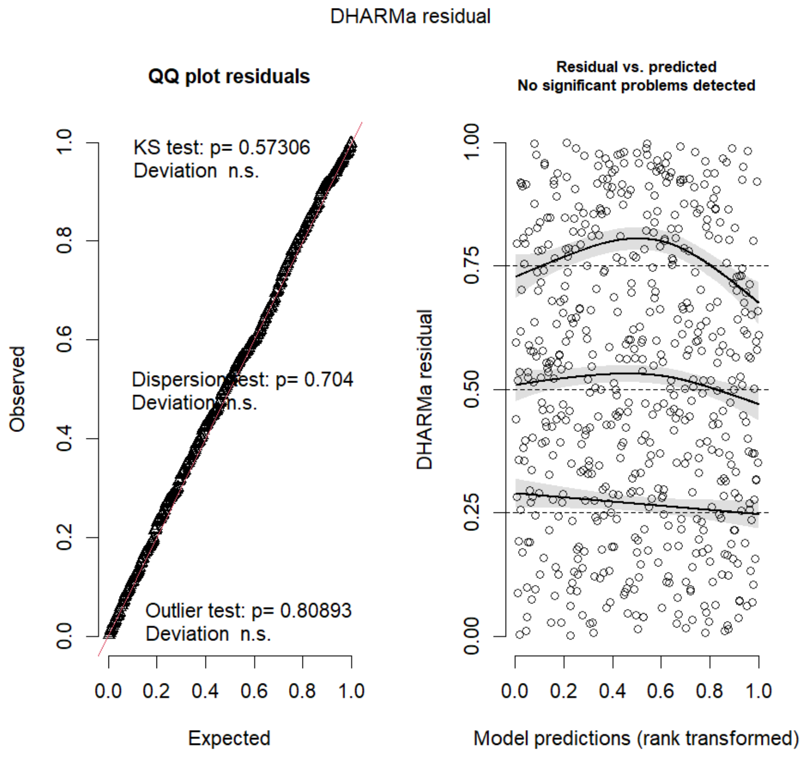

##### Table S28: Analyss of deviance - Type II Wald Chi-square Tests for Model 10

| Term | Chisq | Df | Pr(>Chisq) |
|---|---|---|---|
| Strategy | 0.2065 | 2 | 0.90189 |
| Chose_Diff | 1.6477 | 1 | 0.19927 |
| I(Chose_Diff^2) | 6.1207 | 1 | 0.01336* |

#### Supplement S15: Details for Model 11

##### Figure S16: Diagnostic Plots for Model 11

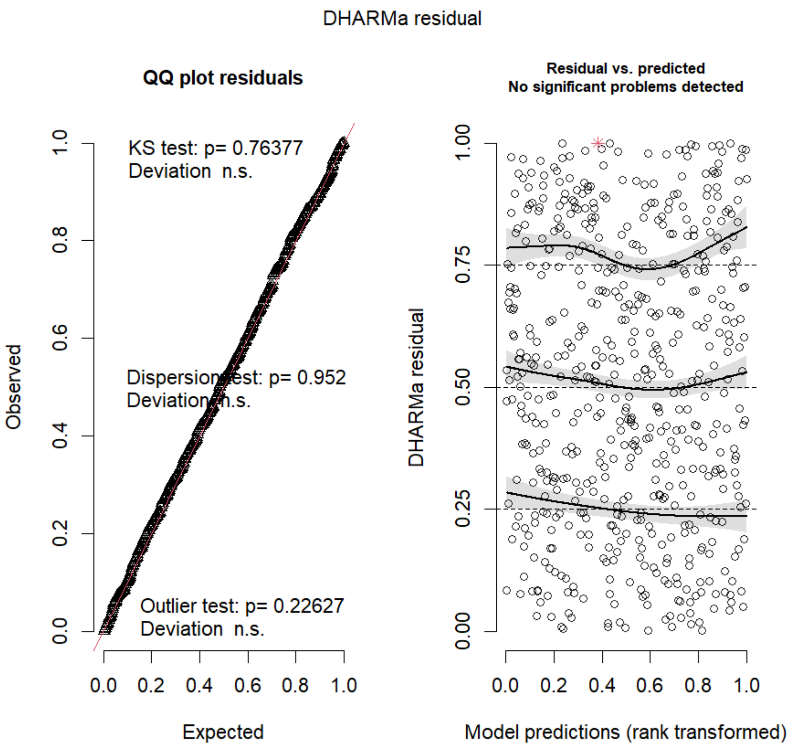

##### Table S29: Analyss of deviance - Type II Wald Chi-square Tests for Model 11

| Term | Chisq | Df | Pr(>Chisq) |
|---|---|---|---|
| Strategy | 1.1813 | 2 | 0.55397 |
| Chose_Diff | 2.8034 | 1 | 0.09406. |
| I(Chose_Diff^2) | 15.4901 | 1 | 8.294e-05*** |

### Supplement S16: Details for Model 12

#### Figure S17: Diagnostic Plots for Model 12

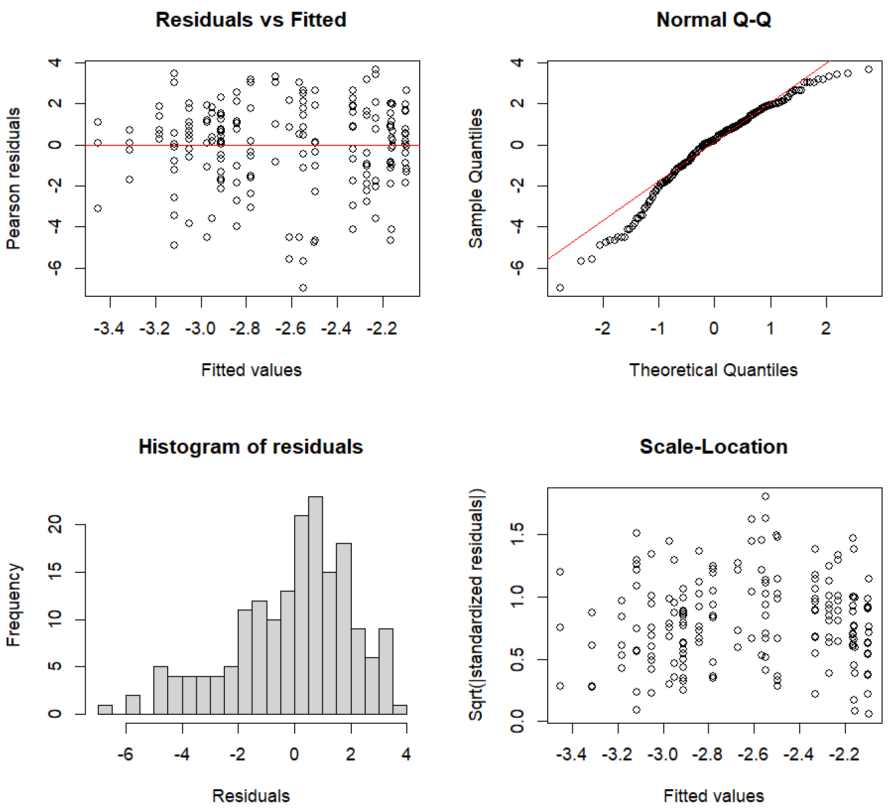

#### Table S30: Analyss of deviance - Type II Wald Chi-square Tests for Model 12

| Term | Chisq | Df | Pr(>Chisq) |
|---|---|---|---|
| Strategy | 11.484 | 2 | 0.003208** |

#### Table S31: First Emmeans Contrast for Model 12

| Contrast | estimate | SE | df | t.ratio | P value |
|---|---|---|---|---|---|
| Average-Fast | 0.339 | 0.124 | 172 | 2.741 | 0.0185* |
| Slow-Average | 0.062 | 0.084 | 172 | -0.737 | 0.7417 |
| Slow-Fast | 0.401 | 0.119 | 172 | -3.379 | 0.0026 |

Results are given on the log (not the response) scale.

P value adjustment: tukey method for comparing a family of 3 estimates

#### Supplement S17: Details for Model 13

##### Figure S18: Diagnostic Plots for Model 13

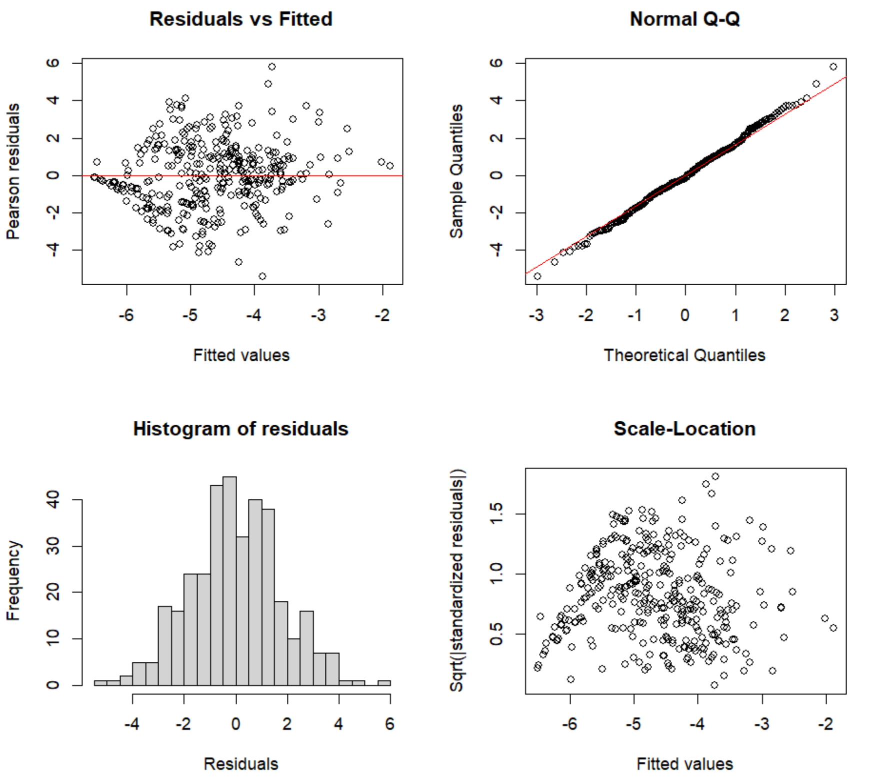

##### Table S32: Analyss of deviance - Type II Wald Chi-square Tests for Model 13

| Term | Chisq | Df | Pr(>Chisq) |
|---|---|---|---|
| Strategy | 5.0034 | 2 | 0.08194. |
| Aggression Type | 25.4417 | 1 | 4.560e-07*** |
| Strategy : Aggression Type | 34.6935 | 2 | 2.927e-08*** |

##### Table S33: First Emmeans Contrast for Model 13

| Contrast | estimate | SE | df | t.ratio | P value |
|---|---|---|---|---|---|
| | | | Type = Attack | | |
| Slow-Average | -0.0143 | 0.171 | 345 | -0.084 | 0.9961 |
| Slow-Fast | 0.3323 | 0.217 | 345 | 0.2760 | 0.2760 |
| Average-Fast | 0.3466 | 0.227 | 345 | 0.2800 | 0.2800 |
| | | | Type = Attacked | | |
| Slow-Average | 0.5835 | 0.171 | 345 | 3.419 | 0.0020 |
| Slow-Fast | 0.4444 | 0.217 | 345 | 2.053 | 0.1014 |
| Average-Fast | -0.1391 | 0.227 | 345 | -0.612 | 0.8134 |

Results are given on the log (not the response) scale.
P value adjustment: tukey method for comparing a family of 3 estimate
